# A cytokinetic ring-driven cell rotation achieves Hertwig’s rule in early development

**DOI:** 10.1101/2023.06.23.546115

**Authors:** Teije C. Middelkoop, Jonas Neipel, Caitlin E. Cornell, Ronald Naumann, Lokesh G. Pimpale, Frank Jülicher, Stephan W. Grill

## Abstract

Cells tend to divide along the direction in which they are longest, as famously stated by Oscar Hertwig in 1884 in his ‘long axis’ rule^1,2^. The orientation of the mitotic spindle determines the cell division axis^3^, and Hertwig’s long axis rule is usually ensured by forces stemming from microtubules^4^. Pulling on the spindle from the cell cortex can give rise to unstable behaviors^5,6^, and we here set out to understand how Hertwig’s long axis rule is realized in early embryonic divisions where cortical pulling forces are prevalent. We focus on early *C. elegans* development, where we compressed embryos to reveal that cortical pulling forces favor an alignment of the spindle with the cell’s short axis. Strikingly, we find that this misalignment is corrected by an actomyosin-based mechanism that rotates the entire cell, including the mitotic spindle. We uncover that myosin-driven contractility in the cytokinetic ring generates inward forces that align it with the short axis, and thereby the spindle with the long axis. A theoretical model together with experiments using slightly compressed mouse zygotes suggest that a constricting cytokinetic ring can ensure Hertwig’s long axis rule in cells that are free to rotate inside a confining structure, thereby generalizing the underlying principle.

## Main

Many cells divide along their long axis, as first described by Oscar Hertwig in 1884^1,2^. This discovery builds on earlier work from Sachs concerning orientations of subsequent cell divisions^7,8^ and is since known as Hertwig’s long axis rule^4,9–15^. Since the mitotic spindle is bisected during cell division for proper chromosome segregation, the axis along which the mitotic spindle is oriented determines the axis of cell division^3^. Hence the orientation of the mitotic spindle determines the axis along which a cell is divided. But how then does the mitotic spindle align with the long axis of a cell, thereby facilitating Hertwig’s rule?

If misaligned, moving the spindle to orient it with the long axis of the cell requires force generation. What types of forces can act upon microtubules and microtubule asters? We distinguish force generation inside the cell, within the cytoplasm, from force generation at the cell cortex. Within the cytoplasm, dynein motors that are attached to cytoplasmic anchors can exert forces on microtubules^16–19^. Assuming that these anchors are present evenly throughout the cytoplasm, the net force generated onto an astral microtubule is proportional to its length. Notably, microtubule asters can position themselves at the center of a cell through length-dependent force generation^9,16,18,20–23^. Moreover, microtubules that grow towards the periphery can continue to do so as they encounter the cell surface, which gives rise to a pushing force^24,25^. Collectively, pushing by astral microtubules can give rise to an elastic restoring force that centers a microtubule aster^26–29^. Finally, active force generation at cell surface anchor sites can exert cortical pulling forces upon astral microtubules and spindle via the dynein-associated protein LIN-5/NuMa^30,31^. Dynein-dependent cortical pulling forces at tricellular junctions in *Drosophila* epithelia, and at retraction fibers in cultured cells, facilitate alignment of the cell division axis with the interphase long axis^10,11^. In addition, the LIN-5/NuMa complex also orchestrates spindle positioning and elongation in the *C. elegans* zygote^32–35^. However, exerting cortical pulling forces onto astral microtubules gives rise to a tug-of-war scenario that can lead to unstable behaviours and even oscillations, which makes it challenging to robustly define positions and orientations^5,6,36^. In principle, all three above mechanisms of microtubule-based force generation can contribute to Hertwig’s long axis rule^9–11,16,18,23,37,38^.

We here set out to investigate how Hertwig’s rule is executed in systems where cortical pulling forces onto astral microtubules are prevalent. To this end, we first study long axis finding in early *C. elegans* blastomeres that divide inside an ellipsoid egg shell^39^. LIN-5/NuMa-dependent cortical pulling forces have been shown to orient mitotic spindles in early blastomere divisions^32–35,40,41^. Since cell polarity cues can override Hertwig’s rule^13,42,43^, we focus on the first symmetric cell division, which is the division of the unpolarized anterior blastomere in the two-cell embryo (Fig. 1A, AB cell). It has previously been shown that, guided by cell-cell contact with the posterior P_1_ cell (Fig. 1A), the AB cell initially sets up the mitotic spindle in the plane orthogonal to the AP axis^44–46^ (the dorsal-ventral left-right, or DV-LR plane, Fig. 1A). The spindle remains in the DV-LR plane up until a cell rotation event at very late stages of cytokinesis^41,44,45,47^ (Fig. S1C right panel). Early embryos are slightly compressed *in utero*^41,48^, such that the AB cell contains a long and a short axis in the DV-LR plane^41^ (Fig. 1A). We first asked whether the AB cell divides along the long axis in this plane. We used spinning-disc confocal microscopy to image the mitotic spindle (GFP::tubulin) and the cell surface (Lifeact::mKate2) of isolated embryos that were mounted under various degrees of compression (see methods). We measured the angle between the mitotic spindle and the long axis of the AB cell in the DV-LR plane (referred to as the spindle angle). While there is no preferred orientation in the absence of compression and for aspect ratios above ∼0.95 (Fig. 1B, C, Movie 1), the mitotic spindle is aligned with the long axis of the cell at the end of anaphase in embryos compressed to aspect ratios below ∼0.95 (spindle angle: 4.4± 20.2 degrees, mean±std throughout the text, n = 40; Fig. 1B, D, Movie 2). Importantly, it was previously reported that the aspect ratio *in utero* is ∼0.86^41^. Altogether, this demonstrates that, when compressed, the dividing AB cell follows Hertwig’s long axis rule. This is so even though the AB spindle is randomly oriented at the onset of cell division and during metaphase (Fig. 1D). Misaligned mitotic spindles then underwent a rotation such that, by the end of anaphase, spindles aligned with the long axis (Fig. 1B, D, Fig. S1A-B, Movie 2). Altogether, Hertwig’s rule is executed in the dividing AB cell via a spindle rotation.

**Fig. 1:**
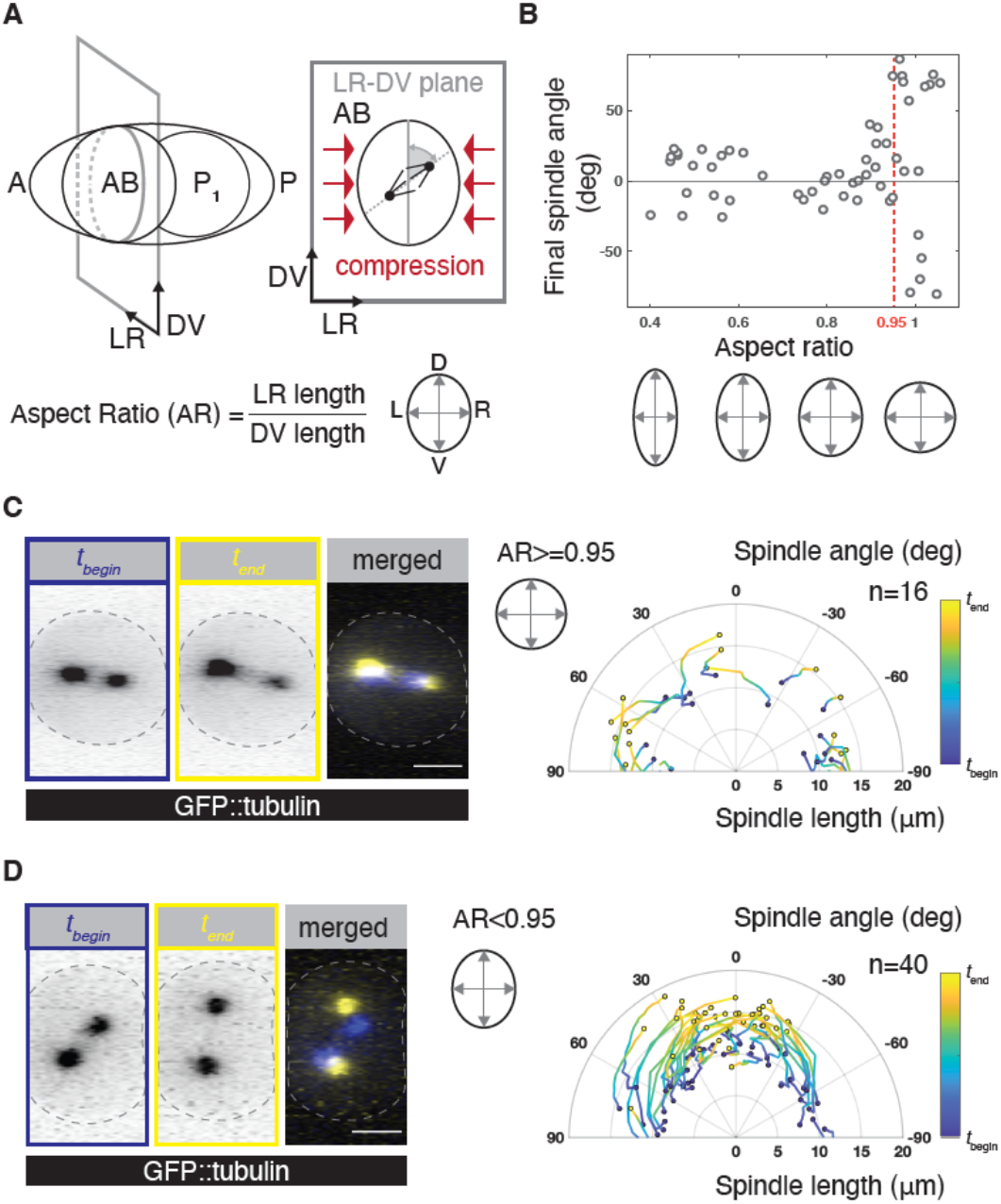
Hertwig’s long axis rule is executed in the AB cell. **A** Left: schematic of a *C. elegans* 2-cell embryo, anterior (A) and posterior (P) are indicated. Right: *In utero*, the embryo is compressed orthogonal to the anteroposterior (AP) axis, such that the AB cell has a long and a short axis in the dorsoventral-left-right (DV-LR) plane. **B** Final angle between the mitotic spindle and the long axis in the DV-LR plane upon different compression strengths. Embryos were dissected and subjected to various degrees of compression, resulting in aspect ratios between 0.4 and 1.1. **C-D** Left panels: Example of an uncompressed (AR>0.95, **C**) and compressed embryo (AR>0.95, **D**), producing GFP-tubulin, at the beginning of anaphase (*t*_*begin*_, blue) and at the end (*t*_*end*_, yellow) viewed in the DV-LR plane. The end point is defined by the starting time point of the cell division skew in the AP-DV plane (Fig. S1C). Dashed lines mark the cell outline, as defined by the Lifeact::mKate2 signal (not shown). Right panels: Time evolution of spindle length (pole-to-pole distance) and angle with the long axis in uncompressed (**C**) and compressed (**D**) embryos, plotted in polar coordinates. Traces represent individual embryos. Time is normalized and subsequently color coded in the same manner throughout the manuscript. For uncompressed embryos, in which there is no long axis, the angle with the imaging plane was reported. Scale bars: 10 um.

We next asked if long axis alignment is driven by LIN-5/NuMa-dependent cortical pulling forces^30^. To address this, we analysed spindle rotation in the AB cell in compressed embryos upon *lin-5/NuMa(RNAi)*^32,49^. Cortical pulling forces were significantly reduced in this condition, as evidenced by a decreased spindle length and reduced transverse spindle pole fluctuations prior to anaphase (Fig. S2). Interestingly, we find that the metaphase spindle in *lin-5/NuMa(RNAi)* embryos was already aligned with the long axis of the cell (initial spindle angle: -4±33.3 degrees, n=12, see supplement), and remained so during anaphase (final spindle angle: 0.2±18.2 degrees, n=12, Fig. 2A, Fig. S1A-B, Movie 3). This is in contrast to unperturbed embryos where the spindle was randomly oriented at metaphase and aligned later during anaphase (Fig. 2B, Movie 4). Together, this indicates that cortical pulling forces counteract long-axis finding during metaphase.

**Fig. 2:**
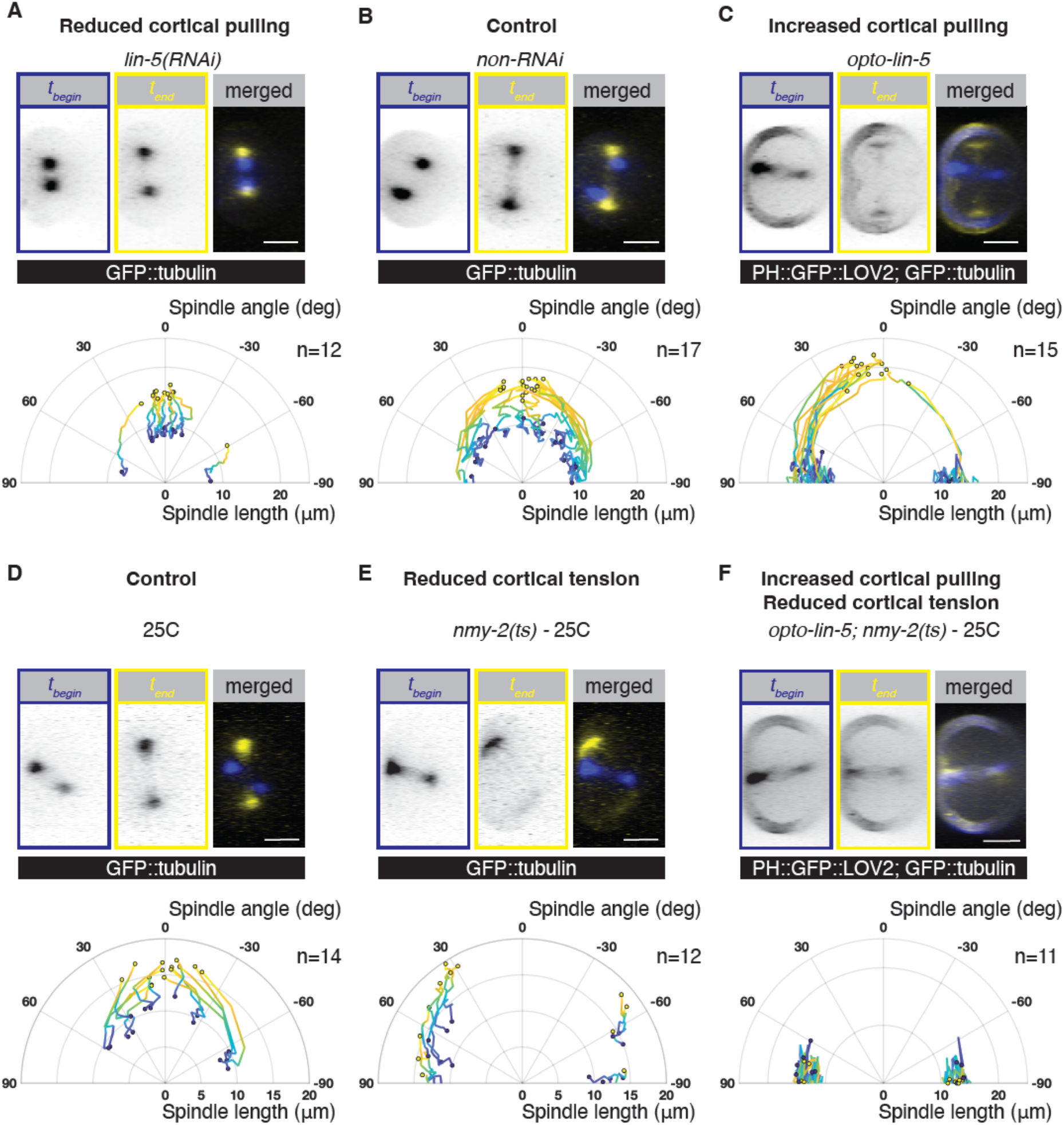
Long axis alignment the AB cell is achieved by an NMY-2 dependent spindle rotation. **A-C** Time evolution of spindle length and angle upon **A** decreased cortical pulling forces - *lin-5(RNAi)*, **B** control - *non-RNAi* and **C** increased cortical pulling forces - *opto-lin-5*. **D-F** Time evolution of spindle length and angle imaged at 25C in **D** control, **E** upon reduced cortical tension - *nmy-2(ts)* and **F** upon combined cortical tension reduction and increased cortical pulling forces - *opto-lin-5; nmy-2(ts)*. Note: the GFP channel in *opto-lin-5* shows GFP::tubulin and PH::GFP::LOV2, which is a membrane localized LOV2 domain necessary to recruit cytoplasmic LIN-5::ePDZ to the cortex (see methods). The control conditions for *opto-lin-5; nmy-2(ts)* imaged at 25C are shown in Fig. S3. n=number of embryos. Temporal color-code as in Fig. 1. Scale bars=10 um.

If Dynein-mediated cortical pulling forces indeed counteract AB spindle long-axis alignment during metaphase, then upregulation of cortical LIN-5/NuMa should lead to metaphase AB spindle short-axis alignment. To test this, we made use of a previously described optogenetic tool, in which cortical LIN-5/NuMa levels are increased upon blue light-illumination (referred to as *opto-lin-5*), thereby increasing pulling forces on astral microtubules^40^. We dissected embryos with minimal light exposure and kept them in the dark until 3-4 min after completion of the first cytokinesis. Thereafter, opto-LIN-5 was globally activated by 3D-imaging of the GFP channel. Cortical pulling forces were significantly elevated in this condition, as evidenced by an increased spindle length and increased transverse spindle pole fluctuations prior to anaphase (Fig. S2, Movie 5). Consistent with our expectation, increasing cortical LIN-5 levels leads to AB metaphase spindles that are aligned with the short axis (initial spindle angle: -85.2±8.6 degrees, n=15, Fig. 2C), a phenotype opposite to that observed in *lin-5/NuMa(RNAi)* (Fig. 2A). However, this initial misalignment is corrected during anaphase via a spindle rotation (final spindle angle: -9±9.5 degrees, n=15; Fig. 2C, Movie 5), thereby ensuring Hertwig’s rule (Fig. S1A-B). Since all our LIN-5/NuMa perturbations affected metaphase, but not anaphase long-axis alignment, we conclude that a LIN-5/NuMa-independent mechanism ensures Hertwig’s rule in this system. Taken together, Dynein-mediated cortical pulling forces favour short-axis alignment of the mitotic spindle during metaphase, which is corrected during anaphase by a LIN-5/NuMa-independent spindle rotation.

We next set out to identify the LIN-5/NuMa-independent mechanism of rotation for achieving long-axis spindle alignment. We note that the actomyosin cortex has also been implicated in Hertwig’s rule albeit with mechanisms that are not well understood^12,30,50–53^. To test if myosin-dependent force generation contributes to Hertwig’s rule in the AB cell, we perturb non-muscle myosin II (hereafter referred to as NMY-2/Myosin), activity by making use of a temperature-sensitive *nmy-2* mutant (*ne3409ts*)^54^. This allele yields functional NMY-2/Myosin at the permissive temperature of 15C, and largely inactive NMY-2/Myosin at the restrictive temperature of 25C^54,55^. Embryos were kept at the permissive temperature until completion of the first cell division, then shifted to the restrictive temperature, followed by analysis of AB spindle dynamics in the DV-LR plane. Acute inactivation of NMY-2/Myosin leads to spindles tending to align with the short axis in the DV-LR plane during metaphase (Fig. 2D-E). This is expected given a reported inhibitory effect of NMY-2/Myosin on cortical pulling forces^56,57^: an acute loss of NMY-2/Myosin activity should therefore lead to increased cortical pulling forces and short axis alignment, as was indeed observed (Fig. 2C). Strikingly however, the rotation of the spindle during anaphase was impaired upon loss of NMY-2/Myosin activity (19.3 ± 12.7 degrees of rotation in *nmy-2(ts)* for embryos where the initial spindle angle was between 30-90 degrees, Movie 7, compared to 47.2±17 in unperturbed control embryos, Movie 6), and the mitotic spindle failed to align with the long axis at the end of anaphase upon acute NMY-2 inactivation (final spindle angle: 68±36 degrees, n=12; Fig. 2E, Fig. S1A-B, Movie 7). This provides evidence that actomyosin drives the rotation during anaphase that ensures Hertwig’s rule in the AB cell. We further tested this possibility by elevating cortical pulling forces in the absence of NMY-2/Myosin activity (*opto*-*lin-5*; *nmy-2(ts)*). As in *opto-lin-5* and *nmy-2(ts)* alone, the metaphase spindle was aligned with the short axis in *opto*-*lin-5*; *nmy-2(ts)* embryos imaged at 25C (initial spindle angle: 85.3±12.6 degrees, n=11; Fig. 2F), but now remained so throughout anaphase, since the NMY-2/Myosin dependent rotation was impaired at the restrictive temperature (final spindle angle: 88.3±6.4 degrees, n=11, Fig. 2F, Fig. S3, Movie 8). Together, we conclude that LIN-5/NuMa-mediated cortical pulling forces and NMY-2/Myosin-driven contractility act antagonistically: LIN-5/NuMa-dependent cortical pulling forces favour spindle short-axis alignment, which is then corrected into spindle long-axis alignment via an NMY-2/Myosin dependent rotation.

How could NMY-2/Myosin drive the rotation that leads to long axis alignment of the spindle? One possibility is that the constriction of the cytokinetic ring could drive a rotation when the overall shape of the cell is fixed by the egg shell^58^. To determine if the cytokinetic ring contributes to long axis alignment in the AB cell, we first investigate the temporal relationship between spindle rotation and cytokinetic ring formation in the AB cell. We perform simultaneous imaging of the actomyosin cortex and the mitotic spindle, and find that spindle rotation is concomitant with the formation and onset of ingression of the cytokinetic ring (Fig. 3A-C, Fig. S4). This analysis also reveals that the ring and the spindle rotate together (Fig. 3D, Fig. S4, Movie 2), suggesting that AB spindle rotation is achieved by rotating the entire AB cell, including all of its intracellular components (Fig. S5A). Furthermore, the AB cell is in direct contact with its neighboring P_1_ cell, which also rotates at the same time (Fig. S5B, Movie 9) indicating that NMY-2/Myosin activity in the AB cell drives both a whole-cell and whole-embryo rotation. Together, this shows that the whole embryo is essentially free to rotate inside the eggshell, consistent with earlier observations^47,48,59,60^.

**Fig. 3:**
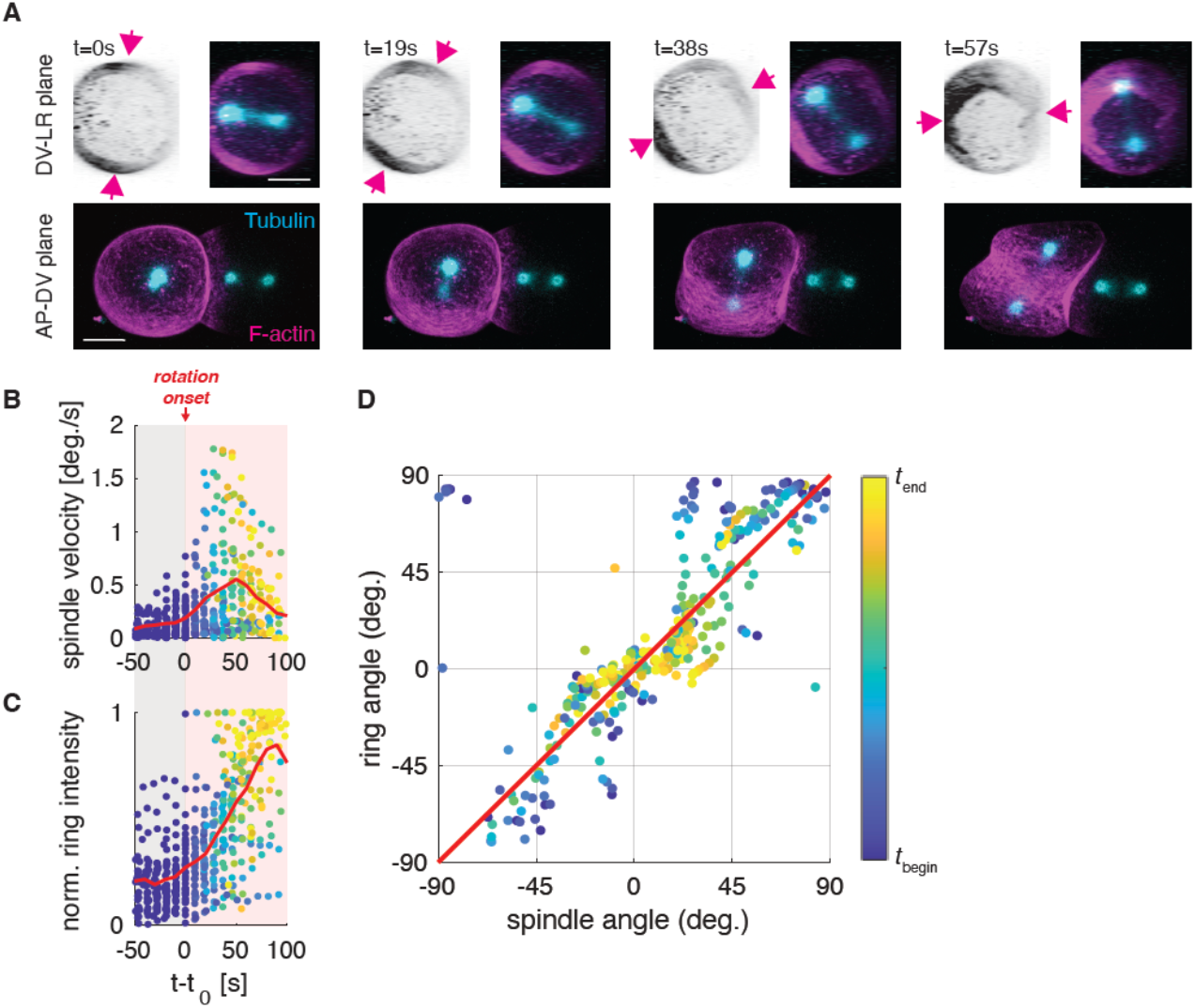
Mitotic spindle and cytokinetic ring rotate together and this coincides with cytokinetic ring formation. **A** Still images of a two-cell embryo producing GFP::tubulin (cyan) and Lifeact::mKate2 (magenta & inverted) viewed in the DV-LR plane (top) and AP-DV plane (bottom). Arrows mark the position of the cytokinetic ring. Scale bar=10 um. **B** Time evolution of the angular velocity of the mitotic spindle and **C** the normalized F-actin intensity in the ring (see Eq. S3 in Supplement). *t*_*0*_ (= *t*_*begin*_) is defined as the rotation onset. Red lines show the mean. As the angular velocity is noisy, data points in **B** represent a moving average with 45s as sliding window. **D** Angle of the spindle and ring during the rotation plotted over time. Note that the angles for spindle and ring are with respect to the long axis and short axis, respectively. The red diagonal is a guide to the eye to indicate the similarity of spindle and ring orientation. Color-coded data points in **B-D** represent individual measurements at respective time points.

How can cell-internal stresses such as the ones generated by NMY-2/Myosin drive a rotation of the entire two-cell embryo? We consider a scenario where the shape of the embryo is fixed by the rigid egg-shell, and where the embryo itself is free to rotate^47,48,59,60^. A rigid body-like rotation of the embryo can result when internal stresses generate a torque acting against frictional forces between the embryo and the egg-shell. This torque arises because the egg-shell exerts forces on the embryo perpendicular to its surface that are required to fix its shape (Fig 4A). The situation during anaphase resembles a purse-string mechanism^58,61–66^: active tension in the cytokinetic ring generates inward forces that will later drive its ingression^67–70^. When the ring of the AB cell is not aligned with the short or long axis of the embryo in the DV-LR plane, the resultant pattern of surface normal forces yields a torque exerted onto the embryo from the eggshell, causing the entire embryo to rotate inside the eggshell (Fig. 4A, supplementary notes). Notably, this rotation of the entire embryo will rotate the cytokinetic ring away from the long axis and align it with the short axis, and thereby align the spindle with the long axis in the DV-LR plane. The same physical argument can be applied to the effects of Dynein-mediated cortical pulling forces by astral microtubules giving rise to inward forces at the cell poles, where astral microtubules are in contact with the cell surface (Fig 4B). However, the torque associated with such forces drives a rotation of the spindle towards short axis alignment (Fig. 4B, supplementary notes). This is consistent with our observations in embryos with elevated cortical pulling and reduced Myosin activity (*opto*-*lin-5*; *nmy-2(ts)*, Fig. 2F). Together, this suggests that for a cell whose shape is constrained but free to rotate inside the egg-shell, Hertwig’s rule is a consequence of cytokinetic ring tension.

**Fig. 4:**
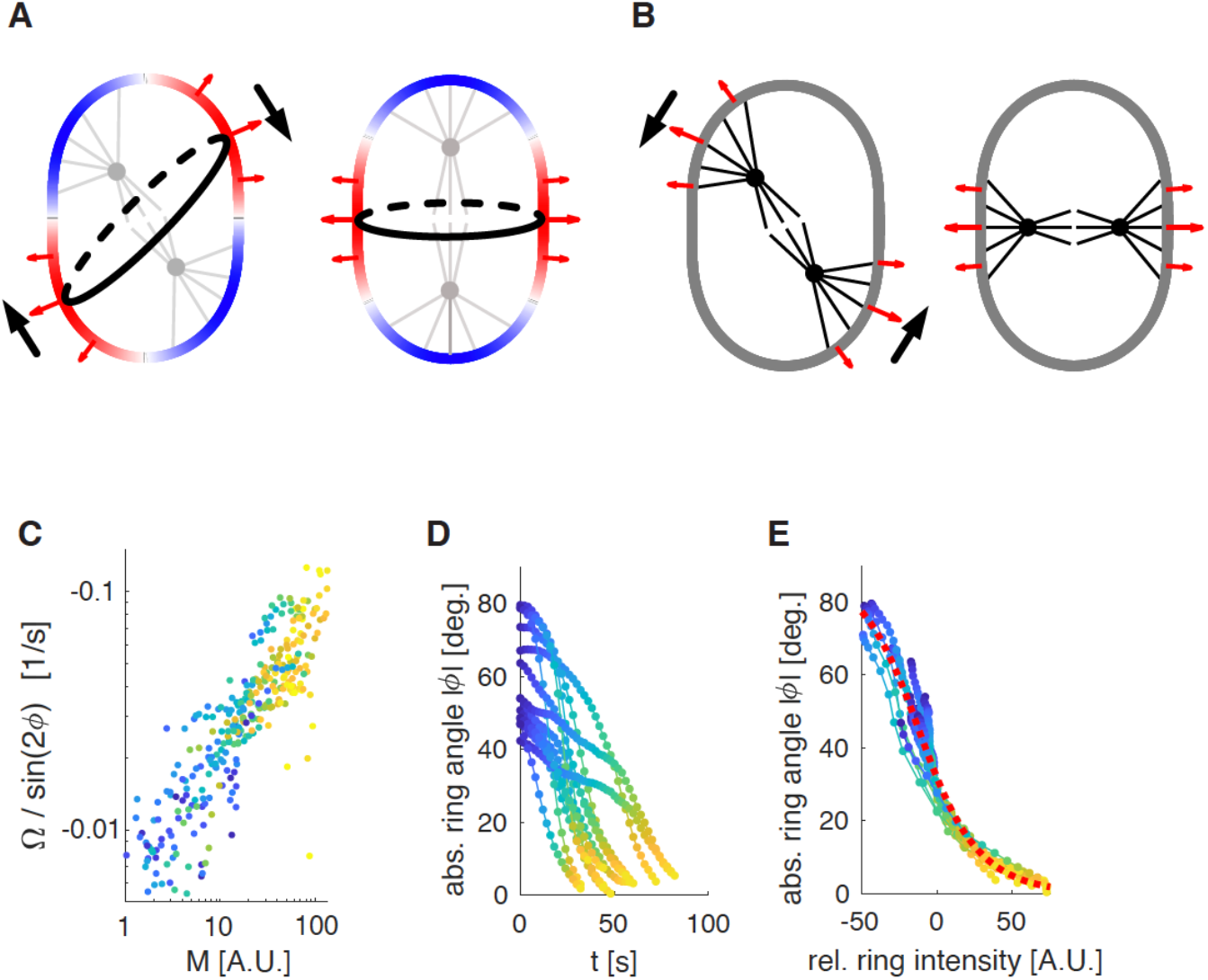
Physical model of long axis alignment driven by the emerging cytokinetic ring. **A:** Schematic of the AB cell in the DV-LR plane: Rigid egg-shell enforces shape of the embryo by exerting forces (red arrows) normal to the embryo surface that balance forces resulting from active surface tension in the cell cortex (colored contour). The cytokinetic ring (black contour) is a region of high tension (red) relative to the cell poles (blue), where astral microtubules (gray) inhibit actomyosin. This results in normal forces driving ring ingression and pole expansion that are balanced by the egg-shell. When the ring is not aligned perpendicular to the long axis (left) a torque arises driving a rotation (black arrows) that aligns the ring perpendicular to the long axis and the spindle with the long axis. **B:** Same as in **A** but for a scenario where normal forces are primarily due to cortical pulling by astral microtubules. In such a scenario the cell poles are pulled are inward, resulting in alignment of the spindle axis with the short axis of the cell. **C-E:** Quantitative analysis of the dynamics of ring angle and NMY-2 concentration in the ring using NMY-2::GFP intensity. We observe a striking correlation (rho=-0.8,p-val<1e-3). **C:** Scatter plot of effective force driving alignment determined from cortical rotation Omega and ring angle phi vs. ring intensity M (Eq. S7). **D:** Trajectories of the absolute ring angle |phi|. **E:** Same trajectories as in **D** but plotted vs. the change in ring intensity relative to the ring intensity at |phi|=30°. Trajectories collapse onto an exponential decay of the tangent of phi as predicted by our model for a linear relationship between active tension and NMY-2 concentration.

To quantitatively test our physical model, we recorded 14 divisions of the AB cell at high time resolution (dt=2s) via spinning-disc microscopy of endogenously labelled NMY-2/Myosin::GFP (Movie 9). Our general theory predicts that the speed of rotation Ω is determined by the angle ϕ between the ring and the short axis and the active tension T generated in the ring according to Ω = −αT *sin* 2ϕ (1), where the coefficient α depends on the aspect ratio of the embryo in the DV-LR plane, the friction between embryo and eggshell, and an effective viscosity of the embryo. Figure 4C reveals that Ω/ *sin* 2ϕ increases monotonously with the intensity of myosin M in the ring, as expected for a linear or saturating relationship between the amount of myosin M and active tension T in the cytokinetic ring. Furthermore, we find that the amount of myosin M increases exponentially with time during ring formation (Fig. S6E), leading to a prediction of the model where the tangent of phi decreases exponentially with increasing M (supp). Indeed, we find that the trajectories of phi collapse onto such a curve when plotting phi vs. M (Fig. 4D-E). Taken together, these results are consistent with a scenario where Herwig’s rule in the AB cell is facilitated by NMY-2/Myosin dependent force generation in the emerging cytokinetic ring, which drives a whole-cell rotation in the case of misalignment.

Given that blastomeres in holoblastic cleavages are not reported to be anchored to the confining shell, we argue that they are often free to rotate. This suggests that the actomyosin-dependent mechanism of long-axis alignment we have identified is general in early blastomere divisions. To evaluate this, we next studied the first division in mouse embryos, in which it has previously been shown that Herwig’s rule applies^14^. We deformed mouse zygotes into an elliptical shape using glass pipettes (Movie 10-12) and found that the cell division axis aligns with the long axis in both mildly and strongly compressed embryos (n=12 embryos, Fig. 5, Fig. S7, Movie 11-12). We next asked if they undergo a whole-cell rotation at the onset of cytokinesis, driven by the actomyosin ring, to align the division axis with the long axis. We found that in mildly compressed zygotes the metaphase spindle orientation was unaffected by compression (n=5 embryos). Importantly, in all cases where the metaphase spindle was misaligned with the long axis of the mildly compressed egg (4 out of 5 embryos), the blastomere underwent a whole-cell rotation at the time of cleavage furrow ingression, thereby aligning the mitotic spindle with the long axis of the cell (Fig. 5, Movie 11). Taken together, experiments with slightly compressed mouse zygotes reveal that also here the entire cell rotates as the cytokinetic ring constricts. These results suggest that the cytokinetic ring can execute Hertwig’s rule in holoblastic blastomere divisions of evolutionary distant species.

**Fig. 5:**
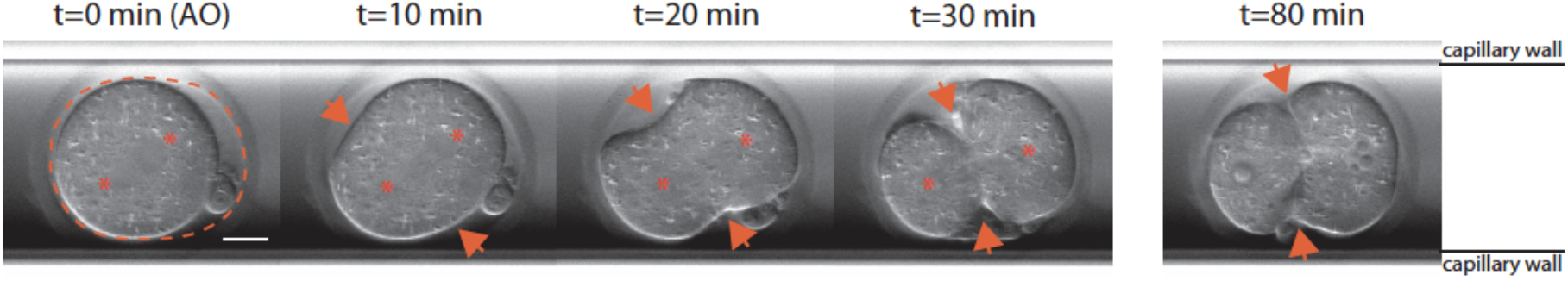
Hertwig’s rule execution in slightly compressed mouse embryos coincides with ring ingression. Differential Interference Contrast (DIC) still images of a mouse zygote undergoing cell division inside a glass capillary. Dashed line indicates the shape of the zona pellucida and asterisks indicate the estimated spindle pole positions. Embryo is slightly compressed, such that the zona pellucida has a long and a short axis. During metaphase, the mitotic spindle is not aligned with the long axis of the zona pellucida but during anaphase it rotates towards the long axis. Arrows mark the cytokinetic ring. Importantly, strongly compressed embryos already have their mitotic spindle aligned during metaphase (Fig. S7). AO = anaphase onset. Scale bar = 20 um.

In summary, we suggest the following mechanism for long axis spindle alignment: in scenarios where cell shapes are constrained and where cells are free to rotate, long axis alignment is a consequence of active tension generation in the cytokinetic ring, which corrects any previous misalignment of the mitotic spindle. Importantly, when cortical pulling forces onto astral microtubules are not dominant, spindles may already align with the long axis of the cell during metaphase, via cytoplasmic pulling forces^9,16,18,20–23^ or via pushing^24,26–29^ (Fig. 2A). Active tension generation in the cytokinetic ring will maintain this alignment. In cases where cortical pulling forces are dominant, spindles first align with the short axis of the cell. However, since the cytokinetic ring establishes in a plane that is orthogonal to the spindle axis, the ring will now not be aligned with the short axes of the cell. Myosin-driven tension in the cytokinetic ring will then drive a whole cell (or whole embryo) rotation that aligns the ring with the short axis, thereby leading to alignment of the spindle with the long axis. Shapes of cells and patterns of surface forces can be more complex than considered here, but the principle of undergoing a rotation to align a pattern of actomyosin-generated surface forces with cell shape remains general (see supplementary notes). We suggest that this actomyosin-dependent mechanism of executing Hertwig’s rule is an important contributor to spindle positioning and cell division orientation during early development.

## Acknowledgements

We thank Britta Schroth-Diez, Romina Piscitello and Riccardo Maraspini from the Light Microscopy Facility of the MPI-CBG, Dresden, for their expert advice and assistance with time-lapse imaging. We are grateful to the 2018 MBL physiology course students and faculty where this work was started. We thank Karin Crell, Friederike Thonwart and Tina Neumann for assistance with *C. elegans* genetics, Alison Kickuth for discussions on blastomere cleavage divisions, Sander van den Heuvel for sharing the *opto-lin-5* strain, Bob Goldstein for sharing *nmy-2(cp8)*, Tony Hyman for sharing the *GFP::tubulin* transgene and Pierre Gönczy for critical comments on the manuscript. Some strains were provided by the CGC, which is funded by NIH Office of Research Infrastructure Programs (P40OD010440). T.C.M. was supported by the Czech Science Foundation (GACR, grant no. 23-07396S). S.W.G. was supported by the Max Planck Society and the European Research Council (grant no. 742712).

## Competing financial interests

The authors declare no competing interests.

## Notes

### Competing Interest Statement

The authors have declared no competing interest.

### Summary of Updates

A typo was found and is now corrected.

