## Supplement for "A cytokinetic ring-driven cell rotation achieves Hertwig’s rule in early development"

#### Supplementary figures

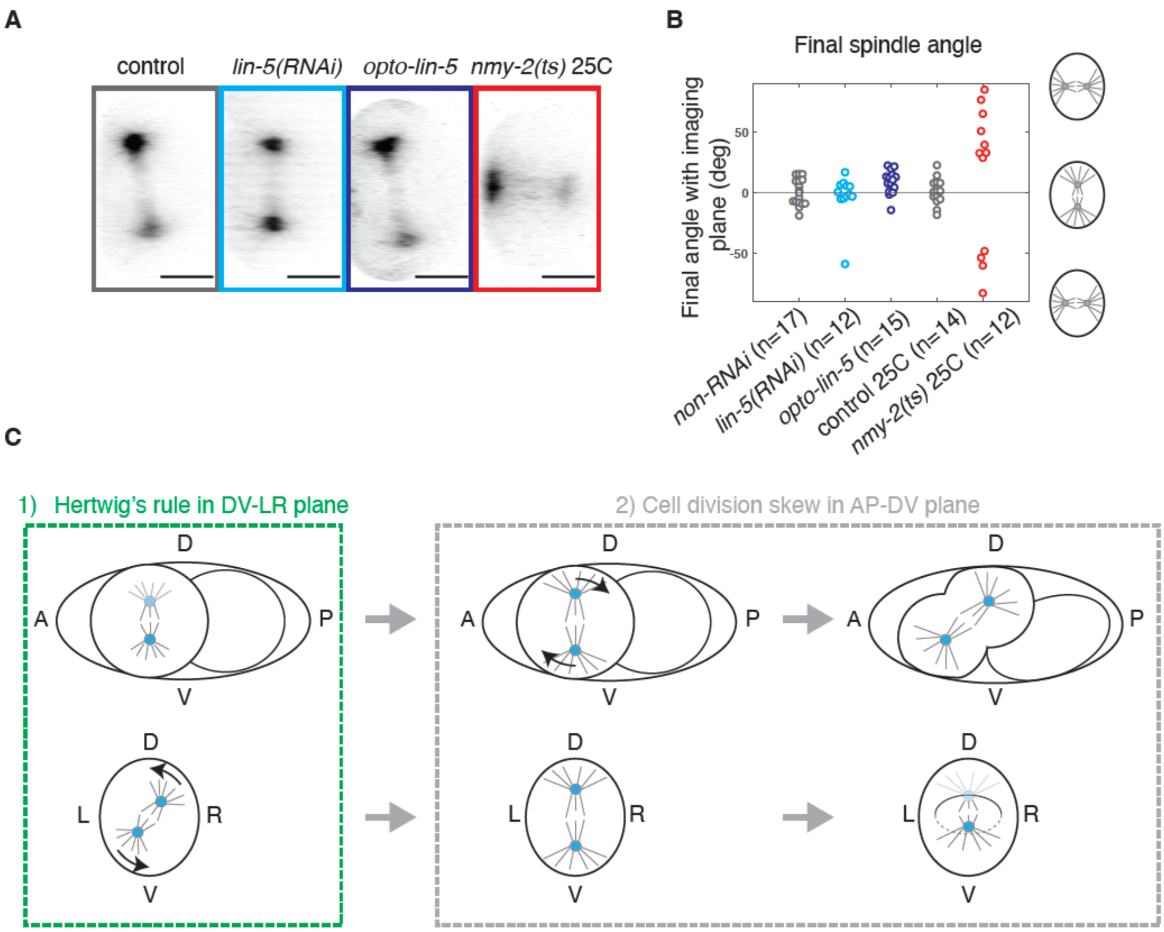

**Fig. S1. Herwig's rule execution in the AB cell upon perturbation of cytoskeletal forces. A** Still images of embryos producing GFP::tubulin in different conditions, defined by the starting time point of the cell division skew in the AP-DV plane. Scale bars: 10 um. **B** Final spindle angle with the long axis upon different perturbations. **C** Schematic of the AB cell division viewed in the AP-DV plane (top) and the DV-LR plane (bottom). Hertwig's rule is executed by a spindle rotation during anaphase in the DV-LR plane (left). This occurs before the cell division skew in the AP-DV plane (right).

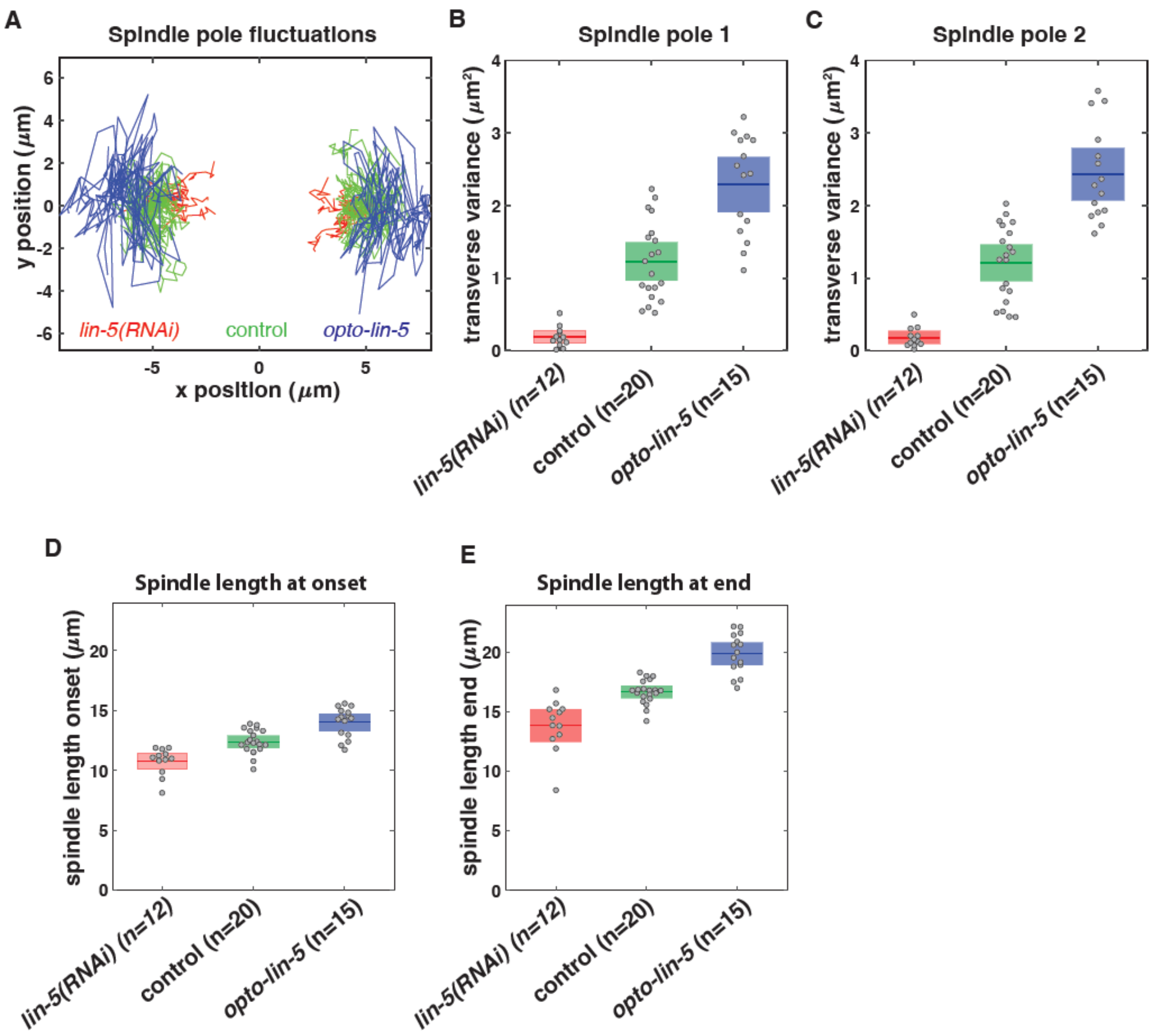

**Fig S2. Modulating cortical LIN-5 levels affects cortical pulling forces on the spindle. A** Spindle pole fluctuations in control (green), *lin-5(RNAi)* (red), and *opto-lin-5* (blue). The spindle poles were tracked in the 15 time points (120 sec) leading up to the onset of rotation, or up to anaphase onset in *lin-5(RNAi)*. Subsequently, for each embryo the spindle pole coordinates were rotated such that the spindle was oriented horizontally, along the x-axis, at the onset of rotation. Figure **A** shows time traces of both spindle pole coordinates in all embryos. **B-C** Variance in y-positions, denoted transverse variance, for each embryo of both spindle poles. Mean over embryos ± 95% confidence intervals are indicated. Increasing levels of cortical LIN-5 lead to more movements in the y-direction, i.e. more transverse fluctuations. **D-E** Spindle length (pole-to-pole distance) at onset and at the end of the rotation. The end point is defined by the starting time point of the cell division skew in the AP-DV plane (Fig. S1C). The onset is defined as 160 seconds prior to the end point.

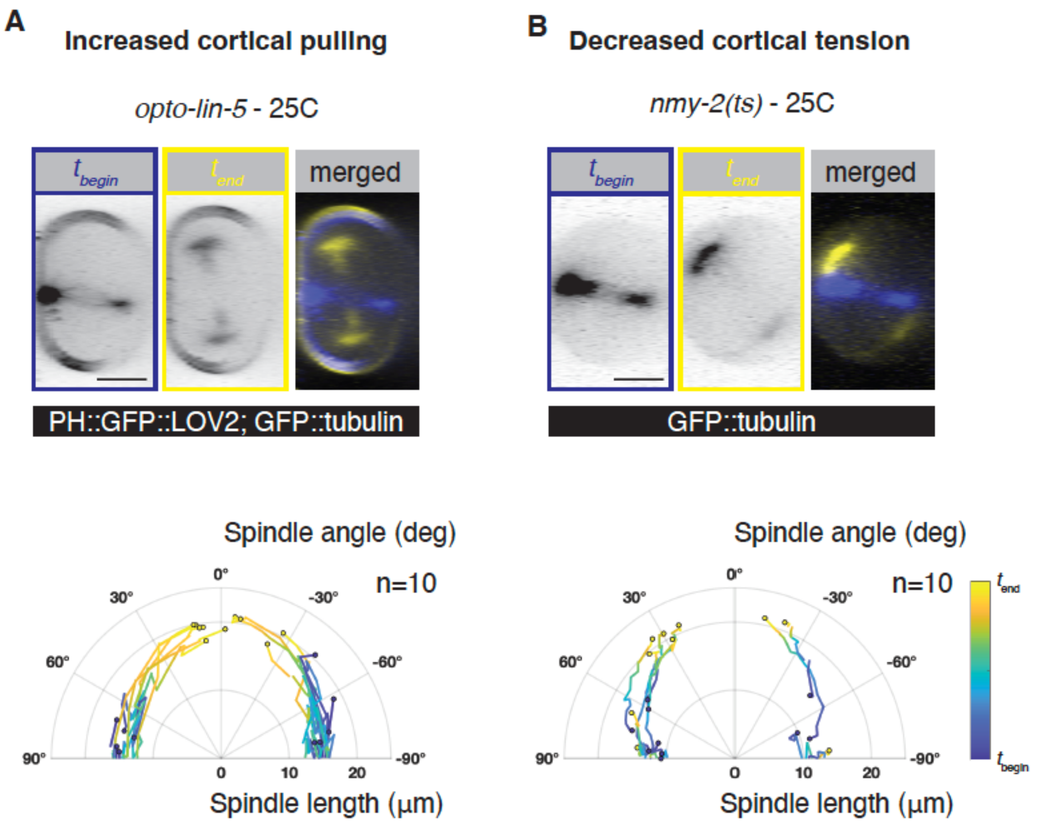

**Fig S3. Controls for the *nmy-2(ts); opto-lin-5* condition in main text.** Spindle positioning in the DV-LR plane of compressed embryos imaged at 25C upon **A** increased cortical pulling on the spindle - *opto-lin-5*, and **B** reduced cortical tension *nmy-2(ts)* strain. Top: still images showing **A** PH::GFP::LOV2; GFP::tubulin and **B** GFP::tubulin at the at the beginning of anaphase ($t_{begin}$, blue) and at the end ($t_{end}$, yellow), viewed in the DV-LR plane. Bottom: time evolution of spindle length (pole-to-pole distance) and angle with the long axis plotted in polar coordinates. Traces represent individual embryos. This supplemental figure provides the controls for the *opto-lin-5; nmy-2(ts)* experiment (main Fig. 2F), which were done in the absence of the *lifeact-mKate2* transgene (unlike main Fig 2E) and at 25C (unlike main Fig. 2C).

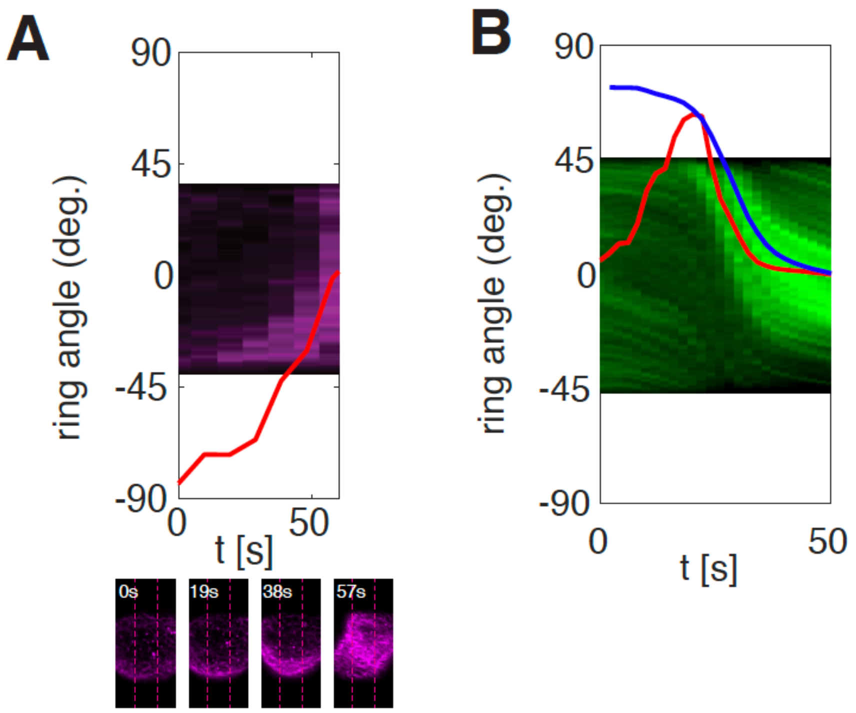

**Fig. S4. Inferring ring angle from cortical intensity measurements. A** Kymograph of Lifeact::mKate2 intensity from the cortical plane of the embryo in Fig. 2 averaged along AP axis in the window indicated with dashed lines in the micrographs below. Each square corresponds to a pixel along the DV axis. The corresponding azimuthal angle was determined from the cortical outline in the DV-LR plane (see Eq. S1 in Supplement). Using linear regression of the fitting function in Eq. S2, the ring angle (red line, Eq. S3) was determined for each time-point. **B** Kymograph as in **A** but for GFP intensity of an embryo producing NMY-2::GFP. Red line corresponds to ring angle determined from kymograph as in **A** and defined in Eq. S3, whereas for the blue line the DV velocity from PIV was used yielding a ring angle as defined in Eq. S4.

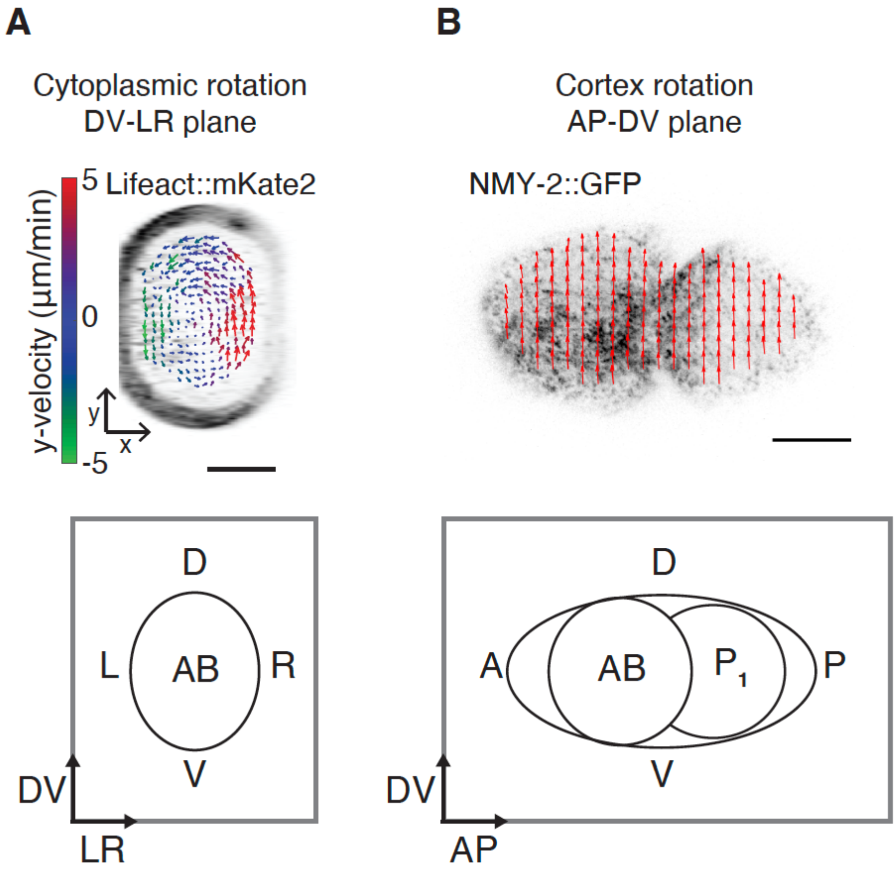

**Fig S5. Both the whole AB cell and the whole embryo rotates during AB cytokinesis. A** Still image of the AB cell of a compressed embryo producing lifeact-mKate2, in the DV-LR plane, undergoing tension-driven rotation. The mean flow field, as measured by particle image velocimetry (PIV) on cytoplasmic F-actin signal, is overlaid. Velocity vectors are color-coded for the component along the future DV axis (y-velocity). Flow filed reveals that the cytoplasm rotates in the same direction as the mitotic spindle and cytokinetic ring. **B** Still image of a compressed two-cell embryo producing NMY-2::GFP, in the AP-DV plane, undergoing tension-driven rotation. The mean flow field as measured by particle image velocimetry (PIV) on cortical NMY-2::GFP signal is overlaid. The AB cell and the neighboring $P_1$ cell rotate in the same direction at the same time. Scale bars = 10 um.

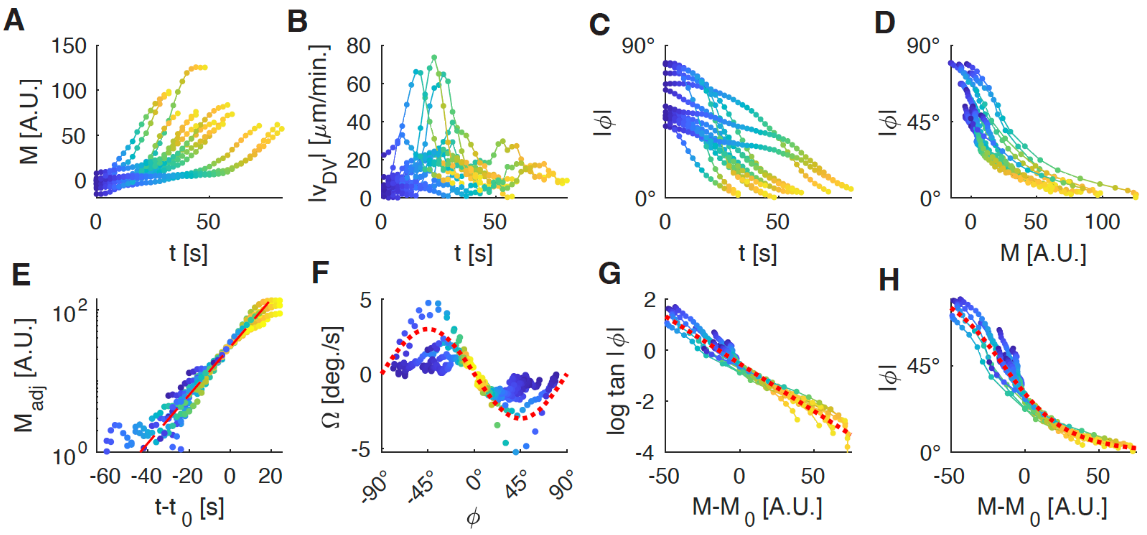

**Fig. S6 Quantification of ring dynamics and comparison to theory using NMY-2-GFP.** All data points are time points from single embryos. Blue and yellow points correspond to first and last time point of rotation. Connected points correspond to subsequent time-points of single embryo. (A) Trajectories of the ring intensity $M_{raw}$ (Eq. S5). (B) Average DV velocity as determined using PIV. (C) Trajectories of the ring angle (Eq. 4) (D) Ring angle plotted as a function of the ring intensity $M_{raw}$. (E) Lin-log plot of the time evolution of the adjusted ring intensity $M$. Allowing for a constant offset (Eq. S7), we find that the ring intensities grow exponentially for early time points with a common growth rate $\lambda = 1/(13s)$. Red dashed line corresponds to exponential fit. (F) Cortical angular velocity $\Omega$ as a function of the ring angle $\phi$. We find that the sign of the angular velocity is always opposite to the sign of the angle such that $\phi = 0°$ corresponds to a stable fixed point and $\phi = 90°$ to an instable fixed point of the dynamics. As expected from a coarse-grained model yielding $\Omega \sim -sin\,2\,\phi$ (Eq. 64 in theory supplement), angular velocities are maximal close to 45°. Red dotted line is $-3\,sin\,2\,\phi$. (G) Plot of $log\,|tan\,\phi\,(t)|$ as a function of the relative ring intensity $M_{raw}(t) - M_{raw}(\phi(t) = 30°)$ reveals that curves from different embryos collapse onto linear curve (red dotted line) with common slope as expected from the coarse-grained model for a linear relationship between Myosin concentration and active surface tension. (H) Same as in (G) but plotting f $|\phi(t)|$. Red dotted curve is given by Eq. S8.

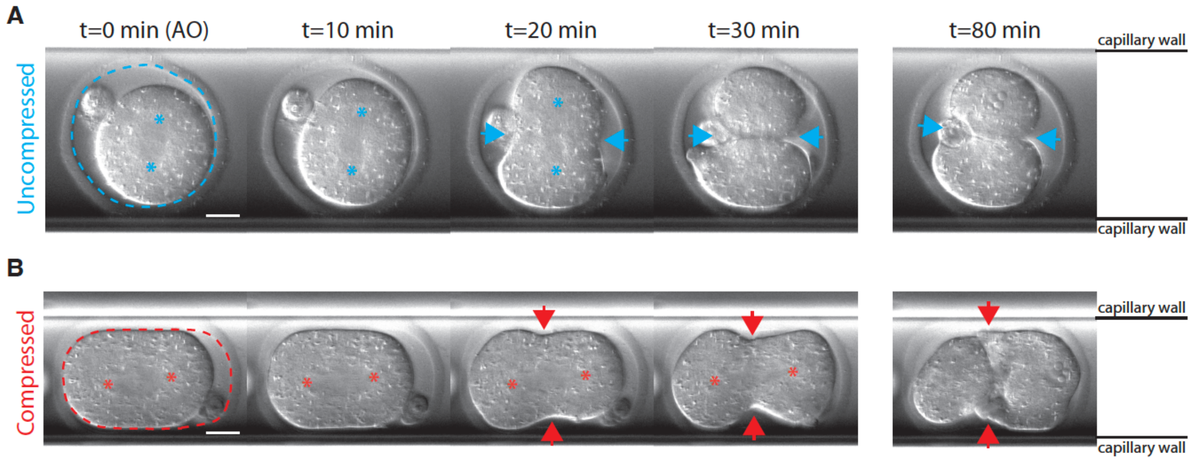

**Fig S7. Mouse zygote division upon different levels of compression.** Differential Interference Contrast (DIC) still images of a non-compressed mouse zygote (n=1 embryo) **A** and a strongly compressed zygote (n=7 embryos) **B** undergoing cell division inside a glass capillary. Dashed line indicates the shape of the zona pellucida and asterisks indicate the estimated spindle pole positions. In strongly compressed zygotes the mitotic spindle already aligns during metaphase, and stays aligned during anaphase. Arrows mark the cytokinetic ring. AO = anaphase onset. Scale bar = 20 um.

**Movie legends**

**Movie 1: Dynamics of the mitotic spindle and actin cortex during AB cell division in an uncompressed embryo**. Spinning disc imaging of an uncompressed embryo producing Lifeact::mKate2 and GFP::tubulin. 3D imaging was performed by making 30 um z-stacks (dz=1um) capturing the entire embryo. Subsequently, maximum intensity projections were made to visualize the DV-LR plane (left panels) and the AP-DV plane (right panels). Due to the lower axial resolution in the z-direction, pixels along the z-axis (LR axis) were interpolated on a grid with 0.1058 um spacing corresponding to the pixel size in the x- and y-axis (AP and DV respectively). Top panels show inverted GFP::tubulin channel alone and bottom panels show the merged Lifeact-mKate2 and GFP::tubulin channels.

**Movie 2: : Dynamics of the mitotic spindle and cortex during AB cell division in a compressed embryo**. Spinning disc imaging of a compressed embryo producing Lifeact::mKate2 and GFP::tubulin. 3D imaging was performed by making 25 um z-stacks (dz=1um) capturing the entire embryo. Rest is as in Movie 1.

**Movie 3: Dynamics of the mitotic spindle during AB cell division in a compressed *lin-5(RNAi)* embryo viewed in the DV-LR plane.** Movie shows GFP::tubulin signal in the AB cell projected onto the DV-LR plane in a *lin-5(RNAi)* embryo producing GFP::tubulin and endogenously labeled LIN-5::ePDZ::mCherry (not visualized).

**Movie 4: Dynamics of the mitotic spindle during AB cell division in a compressed *L4440 control* embryo viewed in the DV-LR plane.** Movie shows GFP::tubulin signal in the AB cell projected onto the DV-LR plane in an *L4440* control embryo producing GFP::tubulin and endogenously labeled LIN-5::ePDZ::mCherry (not visualized).

**Movie 5: Dynamics of the mitotic spindle during AB cell division in a compressed *opto-lin-5* embryo viewed in the DV-LR plane.** Movie shows GFP::tubulin signal in the AB cell projected onto the DV-LR plane in an *opto-lin-5* embryo producing GFP::tubulin, PH::GFP::LOV2 and endogenously labeled LIN-5::ePDZ::mCherry (not visualized).

**Movie 6: Dynamics of the mitotic spindle during AB cell division in a compressed control embryo at 25C viewed in the DV-LR plane.** Movie shows GFP::tubulin signal in the AB cell projected onto the DV-LR plane in a control embryo producing GFP::tubulin and Lifeact::mKate2 (not visualized).

**Movie 7: Dynamics of the mitotic spindle during AB cell division in a compressed *nmy-2(ts)* embryo at 25C viewed in the DV-LR plane.** Movie shows GFP::tubulin signal in the AB cell projected onto the DV-LR plane in an *nmy-2(ts)* embryo producing GFP::tubulin and Lifeact::mKate2 (not visualized).

**Movie 8: Dynamics of the mitotic spindle during AB cell division in a compressed *opto-lin-5; nmy-2(ts)* embryo and its controls at 25C.** Movie panels show GFP::tubulin signal in the AB cell projected onto the DV-LR plane in *opto-lin-5* (left), *nmy-2(ts)* (middle) *and opto-lin-5; nmy-2(ts)* (right) embryos imaged at 25C. Because expression levels of GFP::tubulin varied, the contrast in the movies was differently adjusted.

**Movie 9: Cortical movements in the AB cell during whole embryo rotation viewed in the AP-DV plane.** Top panel: Cortical rotation of an embryo producing endogenously labeled NMY-2::GFP, imaged using high time resolution (dt=2s). Only the cortical surface was imaged in the AP-DV plane and displayed as inverted color. Bottom panel: Same embryo with flow velocity vectors, measured using Particle Image Velocimetry (PIV), overlaid. Both the anterior AB cell and the neighboring $P_1$ cell undergo a similar rotation within the stationary egg shell. Scale bar=10 um.

**Movie 10: Uncompressed mouse zygote undergoing division inside a glass capillary**
Differential Interference Contrast (DIC) movie of a non-compressed mouse zygote undergoing cell division inside a glass capillary. No reorientation of the cell division axis is observed.

**Movie 11: Slightly compressed mouse zygote undergoing division inside a glass capillary**
Differential Interference Contrast (DIC) movie of a slightly compressed mouse zygote undergoing cell division inside a glass capillary. The cell division axis reorients during anaphase in order to ensure Hertwig's rule execution.

**Movie 12: Strongly compressed mouse zygote undergoing division inside a glass capillary**
Differential Interference Contrast (DIC) movie of a strongly compressed mouse zygote undergoing cell division inside a glass capillary. The cell division axis is already aligned along the long axis prior to anaphase and remains aligned.

### Supplementary Notes
Physical theory of an actomyosin-driven Hertwig's rule

**3 Dynamics of axis alignment 11**

#### 1 Hydrodynamic model of the *C. elegans* embryo during AB anaphase

We want to understand how a rotation of the *C. elegans embryo* arises from forces within the cortex and between cortex and spindle. To this end, we model the surface of the embryo as a two-dimensional continuous material with spherical topology. We consider the shape of this surface to be defined by the shape of the rigid egg-shell. Forces within the embryo that are normal to the surface such that they would drive a deformation of this surface are balanced by forces from the egg-shell. In this coarse-grained model, we do not distinguish between cortex and cell membrane and treat the interface between AB and P1 cell as part of the bulk material enclosed by the surface. For simplicity, we will model the surface as a fluid film with homogeneous viscosity. To understand how active forces drive flows in this curved geometry, we will use a covariant formalism as derived in [3].

#### 1.1 Force balance in a curved surface

The surface of the embryo corresponds to a two-dimensional closed manifold with spherical topology. It can be parmetrised parametrised as $\mathbf{X}(s^1, s^2)$, which defines a covariant basis as

$$\mathbf{e}_i = \partial_i \mathbf{X}, \tag{1}$$

where $i \in \{1, 2\}$, and an outward pointing normal vector

$$\mathbf{n} = \mathbf{e}_1 \times \mathbf{e}_2 / |\mathbf{e}_1 \times \mathbf{e}_2|. \tag{2}$$

In section 1.3, an explicit parametrisation for an axisymmetric surface is given with the basis vectors illustrated in Fig. 1a.

The covariant basic defines a metric tensor as

$$g_{ij} = \mathbf{e}_i \cdot \mathbf{e}_j \tag{3}$$

The inverse $g^{ij}$ defines the contravariant basis

$$\mathbf{e}^i = g^{ij} \mathbf{e}_j \tag{4}$$

With this, any vector field $\mathbf{f}$ on the surface, e.g. a force field, can be written as

$$\mathbf{f} = f_n \mathbf{n} + f^i \mathbf{e}_i, \tag{5}$$

where here and in the following we use Einstein sum convention. $f_n$ denotes the component of $\mathbf{f}$ that is normal to the surface, whereas $f^i = \mathbf{e}^i \cdot \mathbf{f}$ denote tangential components.

In a hydrodynamic model, conservation of momentum implies that momentum can only be transported in terms of a flux. The flux of momentum within the surface in direction $i$ is given by $-\mathbf{t}^\mathbf{i}$, where

$$\mathbf{t}^i = t^{ij} \mathbf{e}_j + t^i_n \mathbf{n} \tag{6}$$

is the stress (or tension) tensor of the surface material. Then, momentum conservation yields the force balance equation

$$\nabla_i \mathbf{t}^i = -\mathbf{f}_{\text{in}} - \mathbf{f}_{\text{out}} - \rho \mathbf{a}, \tag{7}$$

Here, $\nabla_i$ denotes the covariant derivative. $\mathbf{f}_{\text{in}} = f^i_{\text{in}} \mathbf{e}_i + f_{\text{in},n} \mathbf{n}$ is the density of the force the inside of the embryo, in particular spindle and cytoplasm, exert on the surface, and $\mathbf{f}_{\text{out}} = f^i_{\text{out}} \mathbf{e}_i + f_{\text{out},n} \mathbf{n}$ is the force density the surrounding material, in particular the egg-shell, exerts on the surface. $\rho \mathbf{a}$ corresponds to an inertia force resulting from a local acceleration $\mathbf{a}$ and a mass density $\rho$. In the following, we will neglect such inertial terms as we will consider a fluid film at low Reynolds number. Expressed in terms of normal and tangential components, the force balance equation then becomes

$$\nabla_j t^{ji} + C^i{}_j t^j_n = -f^i_{\text{out}} - f^i_{\text{in}} \tag{8}$$

$$\nabla_i t^i_n - C_{ij} t^{ij} = -f_{\text{out},n} - f_{\text{in},n}, \tag{9}$$

where $C_{ij} = -\mathbf{n} \cdot \partial_i \partial_j \mathbf{X}$ is the curvature tensor (corresponding to the second fundamental form).

#### 1.2 Constitutive equations

Here, we consider the shape of the surface to be fixed by the egg-shell, i.e. the normal velocity $v_n$ of the fluid film vanishes. Then, Eq. 9 provides a definition for the normal force $f_{\text{out},n}$. For the tangential component of the force from the egg-shell, we use a simple friction force, i.e.

$$f_i^{\text{out}} = -\gamma v_i, \tag{10}$$

where $\gamma$ is a friction coefficient which does not depend on space and time and $v_i$ is the tangential flow field of the embryo surface. We model this surface as a compressible fluid film with active contractility $\chi$ such that the deviatoric tangential stress tensor reads

$$t_{\text{d}}^{ij} = \eta \left( 2\tilde{v}^{ij} + \nu g^{ij} \nabla_k v^k \right) + \chi g^{ij}. \tag{11}$$

Here, $\eta$ is the shear viscosity and $\nu\eta$ is the bulk viscosity with $\nu$ being a dimensionless number. $\tilde{v}^{ij}$ is the shear rate tensor defined as

$$\tilde{v}^{ij} = \frac{1}{2} \left( \nabla^i v^j + \nabla^j v^i - g^{ij} \nabla_k v^k \right). \tag{12}$$

This corresponds to the minimal model of the actomyosin cortex used in [1]. The active isotropic stress $\chi$ corresponds to a density of force dipoles within the surface resulting from the activity of motor molecules, in particular Myosin. For a contractile cortex, $\chi > 0$. In general, the stress tensor contains also equilibrium contributions $\mathbf{t_e^i}$ resulting from the free energy of the fluid film in the absence of activity. The tangential force resulting from the equilibrium stress is given by a Gibbs-Duhem relation [3] and reads

$$\mathbf{e}_j \cdot \nabla_i \mathbf{t}_e^i = -\sum_I c^I \partial_j \mu^I, \tag{13}$$

where $c^I$ and $\mu^I$ are concentration and chemical potential of chemical species $I$ respectively. This corresponds to a pressure resulting from concentration gradients. Here, we consider a regime where exchange with the cytoplasm limits differences in chemical potential such that the resulting force (given in Eq. 13) is small compared to the force $\nabla_i {t_d}^i{}_j$ resulting from viscosity and active stress. Hence, the equilibrium stress does not contribute to the tangential force balance equation which defines the flow field. This allows us to omit the equilibrium stress in the following, where we discuss the flow field of a non-deforming surface. For simplicity we also do not consider deviatoric contributions to the bending moment that would give rise to a normal stress $t_{\text{n}}^i$. Then, we have

$$t^{ij} = t_{\text{d}}^{ij}, \quad t_{\text{n}}^i = 0. \tag{14}$$

The tangential force balance equation (Eq. 8) then reads

$$\gamma v_i = F_i^{\text{visc}} + F_i^{\text{res}}, \tag{15}$$

with

$$F_i^{\text{visc}} := \eta \left( (\nu+1)\partial_i(\nabla_k v^k) + \epsilon^j{}_i \partial_j(\epsilon^{kl}\nabla_k v_l) + 2\kappa v^i \right) \tag{16}$$

$$F_i^{\text{res}} := \partial_i \chi + f_i^{\text{in}}, \tag{17}$$

where $\epsilon_{ij}$ is the antisymmetric Levi-Cevita tensor and $\kappa = \det C_i{}^j$ is the Gaussian curvature. We have used here that the commutator of the covariant derivative on a curved surface can be written as

$$[\nabla_i, \nabla_j]v^k := (\nabla_i \nabla_j - \nabla_j \nabla_i)v^k = \kappa(\delta_i^k g_{jl} - g_{il}\delta_j^k)v^l. \tag{18}$$

We do not specify $\mathbf{f}_{\text{in}}$ at this point, except that it must be due to interactions within the embryo. This means that it can be written in terms of the three-dimensional stress tensor $\sigma_{\alpha\beta}$ of the bulk of the embryo as

$$f_\alpha^{\text{in}} = -\sum_{\beta \in \{x,y,z\}} \sigma_{\alpha\beta} n_\beta, \quad \alpha \in \{x,y,z\}. \tag{19}$$

Here, $n_\beta$ are the cartesian components of the outward pointing normal vector of the surface. We do not consider external forces acting on the bulk of the embryo like gravitation, implying

$$\sum_{\beta \in \{x,y,z\}} \partial_\beta \sigma_{\alpha\beta} = 0. \tag{20}$$

Hence, the net force the embryo exerts on the surface has to vanish, i.e.

$$\int_{\mathcal{S}} \mathrm{d}S \mathbf{f}_{\text{in}} = -\int_{\mathcal{V}} \mathrm{d}V \partial_\beta \sigma_{\alpha\beta} = 0, \tag{21}$$

where $\mathcal{V}$ is the bulk volume of the embryo. We also do not consider external torques, e.g. from a magnetic field. Therefore, angular momentum conservation implies that $\sigma_{\alpha\beta}$ is symmetric and that the net torque the embryo exerts on the surface has to vanish, i.e.

$$\int_{\mathcal{S}} \mathrm{d}S \mathbf{X} \times \mathbf{f}_{\text{in}} = 0. \tag{22}$$

We also do not consider external forces acting on the surface

$$\int_{\mathcal{S}} \mathrm{d}S \mathbf{f}_{\text{out}} = 0 = \int_{\mathcal{S}} \mathrm{d}S \mathbf{X} \times \mathbf{f}_{\text{out}} \tag{23}$$

This turns out to be crucial for understanding the compression-triggered rotation.

##### 1.3 Geometry of an almost axisymmetric surface

In uncompressed conditions, the shape of the egg-shell and, hence, the embryo is almost axi-symmetric. Hence, the shape of the embryo surface can be written as

$$\mathbf{X}(\theta, s) = (\rho(s)\cos\theta,\ \rho(s)\sin\theta,\ z(s))^T = \rho(s)\boldsymbol{\rho} + z(s)\mathbf{z}. \tag{24}$$

Here, $s$ is the arc-length corresponding to the AP direction on the cortex and $\theta$ is the azimuthal angle. $\rho$ is the distance from the $z$ axis connecting the AP poles. In the arc-length parametrisation $\rho$ and $z$ define an angle $\psi(s)$ via $(\cos\psi, \sin\psi) = (\rho'(s), z'(s))$ (see Fig. 1a). With this, the local basis vectors are given by

$$\mathbf{e}_\theta = \rho\boldsymbol{\theta}, \quad \mathbf{e}_s = \cos\psi\boldsymbol{\rho} + \sin\psi\mathbf{z}, \tag{25}$$

where $\boldsymbol{\theta}, \boldsymbol{\rho}, \mathbf{z}$ are normalized vectors. The curvature tensor is given by

$$C_s{}^s = \psi'(s), \quad C_\theta{}^\theta = \frac{\sin\psi}{\rho}. \tag{26}$$

For small deviations from axisymmetry, the shape can be written as

$$\mathbf{X}' = \mathbf{X} + \delta X_n \mathbf{n}_0, \tag{27}$$

where $\mathbf{n}$ is the outward pointing normal vector given by

$$\mathbf{n} = \sin\psi\boldsymbol{\rho} - \cos\psi\mathbf{z}. \tag{28}$$

Upon such a deformation, the basis vectors change in first order of the deformation as

$$\delta\mathbf{e}^i = -\,\delta X_n C^i{}_j \mathbf{e}^j + g^{ij}(\partial_j \delta X_n)\mathbf{n}, \tag{29}$$

$$\delta\mathbf{n} = -\,(\partial_i \delta X_\mathrm{n})\mathbf{e}^i = -\rho(\partial_\theta \delta X_n)\boldsymbol{\theta} - (\partial_s \delta X_n)(\cos\psi\boldsymbol{\rho} + \sin\psi\mathbf{z}). \tag{30}$$

### 2 Rotation triggered by compression

#### 2.1 Compression triggers rotation as a consequence of torque balance

Experimentally, we find that the compression induced rotation during anaphase of the AB cell resembles the rotation of a rigid body around the AP axis. In the absence of compression, no such rotation is observed. This can be understood as a consequence of torque balance (Eq. 23). To this end, let us write the velocity field as

$$\mathbf{v} = \Omega\rho\boldsymbol{\theta} + \mathbf{v}_\mathrm{res}, \tag{31}$$

where

$$\Omega := \frac{1}{\Theta_{zz}}\mathbf{z} \cdot \int_{\mathcal{S}} \mathrm{d}S\mathbf{X} \times \mathbf{v}, \quad \Theta_{zz} = \int_{\mathcal{S}} \mathrm{d}S\rho^2, \tag{32}$$

$$\mathbf{v}_\mathrm{res} := \mathbf{v} - \Omega\rho\boldsymbol{\theta}. \tag{33}$$

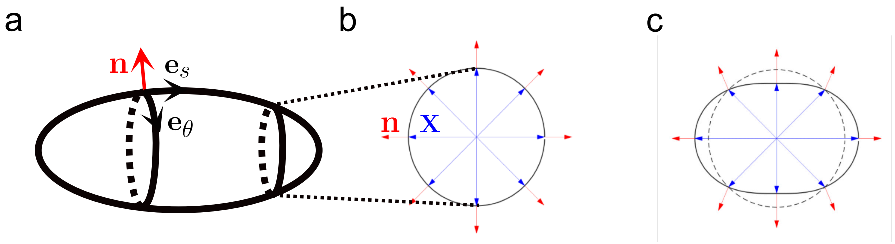

Figure 1: Geometry of an almost axisymmetric surface. (a) axisymmetric surface with tangential vectors $\mathbf{e}_s$, $\mathbf{e}_\theta$ and normal vector $\mathbf{n}$. (b) Cross-section ($xy$ plane) of axisymmetric surface is circle such that position vector $\mathbf{X}$ is parallel to $\mathbf{n}$ in this plane. Hence, $\mathbf{z}\cdot(\mathbf{X}\times\mathbf{n})=0$. (c) Upon non-axisymmetric deformation of the surface, e.g. due to compression of the embryo, the cross-section becomes non-circular and $\mathbf{z}\cdot(\mathbf{X}\times\mathbf{n})\neq 0$ in general.

$\Omega$ corresponds to the component of the flow field that defines a (rigid body) rotation around the $z$ axis. The friction force acting against this rotation yields a torque given by

$$T_{\text{fric}} = -\gamma\mathbf{z}\cdot\int_{\mathcal{S}} \mathrm{d}S\mathbf{X}\times\mathbf{v} = -\gamma\Theta_{zz}\Omega. \tag{34}$$

However angular momentum dictates that the net torque the egg-shell exerts on the embryo surface has to be zero:

$$0 = T_{\text{out}} = \mathbf{z}\cdot\int \mathrm{d}S(\mathbf{X}\times\mathbf{f}_{\text{out}}) = T_{\text{fric}} + T_{\text{n}}. \tag{35}$$

The torque due to normal forces

$$T_{\text{n}} = \mathbf{z}\cdot\int_{\mathcal{S}} \mathrm{d}S\mathbf{X}\times(f_{\text{out,n}}\mathbf{n}) \tag{36}$$

vanishes for an axisymmetric surface, because

$$\mathbf{z}\cdot(\mathbf{X}\times\mathbf{n}) = 0. \tag{37}$$

This can be understood from a cross-section perpendicular to $\mathbf{z}$ which is a circle for an axisymmetric surface (Fig. 1). Hence, the torque from friction force $T_{\text{fric}} = -T_{\text{n}}$ has to vanish. Thus, the rigid body rotation $\Omega$ of an axisymmetric surface has to vanish in the absence of friction gradients or external torques.

Upon compression, a rapid rotation of the embryo is observed during AB anaphase. To understand the effect of compression, we consider a small deformation $\delta X_n << R$ of the embryo surface relative to some axisymmetric reference shape with $R$ being the average diameter of the embryo in the LR-DV plane. With this we show in the following that the compression-triggered rotation can be understood from the balance of torques acting on the embryo surface. In section 2.2, we explicitly calculate how the flow field on the surface of a spherical cell changes upon a small deformation and find that the compression-triggered rotation dominates in the regime of vanishing friction.

On the slightly deformed axisymmetric surface, we write the change in the flow field with respect to a reference flow field $\mathbf{v}_0$ on the axisymmetric surface as

$$\delta\mathbf{v} = \delta\mathbf{v}_{\text{res}} + \delta\Omega\rho\boldsymbol{\theta}, \quad \delta\Omega := \frac{1}{\Theta_{zz}} \int_{\mathcal{S}_0} \mathrm{d}S\rho\boldsymbol{\theta}\cdot\delta\mathbf{v}. \tag{38}$$

With this we can write the torque balance equation (Eq. 23) in linear order of $\delta X_n$ as

$$\delta T_{\text{n}} = -\delta T_{\text{fric,geo}} + \gamma\Theta_{zz}\delta\Omega, \tag{39}$$

where

$$\begin{aligned}\delta T_{\text{n}} =& \mathbf{z}\cdot\int_{\mathcal{S}} \mathrm{d}S\mathbf{X}\times(f_{n,0}^{\text{out}}\delta\mathbf{n}) = -\int_{\mathcal{S}} \mathrm{d}S f_{n,0}^{\text{out}}\partial_\theta\delta X_n \\ =& \int_{\mathcal{S}} \mathrm{d}S(f_{n,0}^{\text{in}} - C_{ij}t_0^{ij})\partial_\theta\delta X_n \qquad (40)\\ \delta T_{\text{fric,geo}} =& -\mathbf{z}\cdot\gamma\int_{\mathcal{S}} \mathrm{d}S\left(\delta\mathbf{X}\times\mathbf{v}_0 + \frac{\delta\sqrt{g}}{\sqrt{g}}\mathbf{X}\times\mathbf{v}_0\right) \\ =& -\gamma\int_{\mathcal{S}} \mathrm{d}S(\rho\psi' + 2\sin\psi)\delta X_n\boldsymbol{\theta}\cdot\mathbf{v}_0. \qquad (41)\end{aligned}$$

Here, $\delta T_{\text{n}}$ is a torque that results from egg-shell normal forces balancing embryo-internal stresses in the presence of a non-axisymmetric deformation $\delta X_n$, as the azimuthal gradient of the deformation $\partial_\theta\delta X_n$ yields a normal vector that is no longer parallel to $\mathbf{X}$ in the LR-DV plane (see Eq. 30 and Fig. 1c). It corresponds to an azimuthal misalignment of the pattern of normal forces $f_{n,0}^{\text{out}}$ and the geometry defined by $\delta X_n$. This can be seen by expanding $f_{n,0}^{\text{out}}$ and $\delta X_n$ as a Fourier series

$$f_{n,0}^{\text{out}}(\theta, s) = \sum_{k\in\mathbb{N}} f_k(s)\cos[k(\theta - \varphi_k^f(s))] \tag{42}$$

$$\delta X_n(\theta, s) = \sum_{k\in\mathbb{N}} d_k(s)\cos[k(\theta - \varphi_k^X(s))], \tag{43}$$

yielding

$$\delta T_{\text{n}} = \pi\sum_{k\in\mathbb{N}} \int_0^L \mathrm{d}s\rho f_k d_k \sin[k(\varphi_k^f - \varphi_k^X)], \tag{44}$$

where we observe that $\delta T_{\text{n}}$ arises from an azimuthal misalignment $\varphi_k^f - \varphi_k^X$ of the pattern of normal forces and the non-axisymmetric geometry.

This torque can trigger a rotation driving an alignment of force pattern and geometry. For simplicity and motivated by the experimentally observed speed of the compression-triggered rotation, we consider here a regime of small friction with the egg-shell, i.e. a large hydrodynamic length $\eta/\gamma >> R^2$. Then, the friction forces $\gamma \mathbf{v}_0$ resulting from the cortical flow towards the cytokinetic ring are small compared to the normal forces $f^{\rm in}_{n,0} - C_{ij} t_0^{ij}$ that drive ingression of the cyokinetic ring and expansion of the cell poles. In this regime, misalignment of normal forces and geometry result in a torque $T_n$ that is large compared to the torque from friction forces $T_{\rm fric,geo}$, such that Eq. 39 simplifies to

$$\delta\Omega = \frac{1}{\gamma\Theta_{zz}}\delta T_{\rm n} = \frac{\pi}{\gamma\Theta_{zz}}\sum_{k\in\mathbb{N}}\int_0^L {\rm d}s \rho f_k d_k \sin[k(\varphi_k^f - \varphi_k^X)]. \tag{45}$$

This means that an misalignment $\varphi_k^f - \varphi_k^X$ between normal forces and geometry triggers a rotation, whenever friction forces with the egg-shell are small compared to the normal forces triggering the rotation. In the regime of small friction we are considering here, the rotation will be fast, i.e. $\delta\Omega\rho >> \langle|\mathbf{v}_0 \delta X_n/\rho|\rangle$. However, a rigid body rotation does not contribute to the normal force $C_{ij}t_0^{ij}$ resulting from viscous forces in the axisymmetric surface. Hence, Eq. 45 remains valid in first order of $\delta X_n$ up to $\delta\Omega\rho$ being of order $\mathbf{v}_0$ at which point viscous forces will limit the rotation. In the AB cell, the compression-independent flow speed prior to chiral flows is about $7\mu$m/min. [2]. We find that the speed of the compression-triggered rotation is about $0.5 - 2$deg./s (Fig. 3) for aspect ratios $AR < 0.95$, corresponding to a cortical flow velocity of $10-30\mu$m/min. (see also Fig. S6B for direct measurements of cortical flow). This suggests that the linear approximation is valid up to $AR \sim 0.95$, corresponding to $\delta X_n \sim \delta\rho \sim 0.3\mu$m, where $\delta\Omega\rho_0 \sim |v_0| = 7\mu$m/min. and hence $\gamma \sim (\eta/\rho^2)(\delta\rho/\rho_0)$. This yields an estimate for the hydrodynamic length,

$$l_h = \eta/\gamma \sim 100\mu\text{m}, \tag{46}$$

which is consistent with the observation that the entire embryo of length $\sim 50\mu$m rotates like a rigid body.

In the absence of external torques and for small deformations $\delta X_n$, the bulk of the embryo will move along with this rigid-body-like rotation of the embryo surface. As the pattern defining the normal forces rotates with the embryo, the rotation (Eq. 45) results in an alignment

$$\partial_t \varphi_k^f \sim \delta\Omega \tag{47}$$

towards

$$\varphi_k^f - \varphi_k^X = 0 + 2nk\pi, \quad n \in \mathbb{N}, \tag{48}$$

for $d_k f_k < 0$. In other words, the deformation-triggered rotation azimuthally aligns patches that pull (push) on the egg-shell such that $f_{\rm n}^{\rm out} > 0$ ($f_{\rm n}^{\rm out} < 0$ with points in the geometry that are deformed inward (outward) relative to the axisymmetric surface, i.e. $\delta X_n < 0$ ($\delta X_n > 0$).

A compression defining a long and a short axis in the LR-DV plane corresponds to a deformation

$$\delta X_n \approx d_2(s)\cos(2(\theta - \varphi_2^X)). \tag{49}$$

For $d_2(s) > 0$, $\varphi_2^X$ is the azimuthal angle of the long axis. The compression-triggered rotation aligns this long axis with the axis of the azimuthal pattern of normal forces, i.e. the $k = 2$ component of $f_\mathrm{n}^\mathrm{out}$ (Eq. 42). For the AB cell dividing in the LR-DV plane with ingressing ring and expanding poles, $\varphi_2^f$ corresponds to the angle of the spindle axis for $f_2 < 0$. Hence, the compression-triggered rotation aligns the spindle axis with the long axis of the cell in the LR-DV plane.

This suggests that Hertwig's rule, i.e. cells dividing along the long axis, is a robust consequence of torque balance, whenever the surface of the cell is free to rotate and the surrounding resists ingression of the ring and/or expansion of the cell poles. Torque balance also provides an explanation for the spindle orientation in embryos with inhibited actomyosin contractility and enhanced astral pulling forces. In these embryos, where we expect the pulling forces at the poles to dominate over ingressing forces at the ring, the spindle aligns with the short axis in the LR-DV plane. We want to stress that this result only depends on the normal forces exerted on the surface in the absence of the deformation $\delta X_n$. Hence it does not require detailed knowledge about the nature of spindle-cortex interactions. It only depends on whether astral microtubules are pushing or pulling at the cortex. Furthermore, we note that this argument is also valid for a scenario where spindle anchors move through the cortex such that the cortex acts as an effectively rigid surface. Given the high viscosity of the cortex and the limited number of anchors, such a scenario seems likely.

#### 2.2 Deformation-triggered rotation of a spherical cell

In the following, we give explicit results for the change in cortical flow that results from statically deforming a spherical cell. This includes the deformation-triggered rotation.

We parametrize the surface as

$$\mathbf{X}(\theta, \phi) = (R + \delta R(\theta, \phi))\mathbf{r}(\theta, \phi), \tag{50}$$

where

$$\mathbf{r}(\theta, \phi) = (\cos\phi\sin\theta, \sin\phi\sin\theta, \cos\theta)^T. \tag{51}$$

$\delta R(\theta, \phi)$ corresponds to a normal deformation of the spherical surface, corresponding to $\delta X_n$ in the previous section. To compute the flow field we use a Hodge decomposition:

$$v_i = \partial_i A + \epsilon^{ji}\partial_j B, \tag{52}$$

where $A$ is a scalar field corresponding to the irrotational component and $B$ is a pseudoscalar corresponding to the rotational component of the tangential flow field $v_i$. We expand these (pseudo-)scalar fields as well as the deformation $\delta R$ and the contractility $\chi$ in terms of scalar spherical harmonics $Y_{lm}$:

$$
\begin{aligned}
A(\theta,\phi) &= \sum_{l=1}^{\infty}\sum_{m=-l}^{l} A_{lm}Y_{lm}(\theta,\phi), \quad B = \sum_{l=1}^{\infty}\sum_{m=-l}^{l} B_{lm}Y_{lm} \\
\delta R_{lm} &= \sum_{l=0}^{\infty}\sum_{m=-l}^{l} \delta R_{lm}Y_{lm}, \quad \chi_{lm} = \sum_{l=0}^{\infty}\sum_{m=-l}^{l} \chi_{lm}Y_{lm}
\end{aligned} \tag{53}
$$

The flow field for $\delta R_{lm} = 0$ is given by

$$
A^0_{lm} = \frac{\chi_{lm}/\eta}{1/l_h^2 + \left[(\nu+1)l(l+1) - 2\right]/R^2}, \quad B^0_{lm} = 0, \tag{54}
$$

where the hydrodynamic length is defined as $l_h = \sqrt{\eta/\gamma}$ For details of the calculation see [1], where also the effect of an enclosed Stokes fluid is considered. For $\delta R \neq 0$, the flow field becomes

$$
A_{lm} = A^0_{lm} + \delta A_{lm}, \quad B_{lm} = \delta B_{lm}, \tag{55}
$$

where

$$
\delta A_{lm} = \frac{1}{1/l_h^2 + \left[(\nu+1)l(l+1) - 2\right]/R^2}\frac{S_{lm} + (-1)^m \bar{S}_{lm}}{2\eta\sqrt{l(l+1)}} \tag{56}
$$

$$
\delta B_{lm} = \frac{-i}{1/l_h^2 + \left[l(l+1) - 2\right]/R^2}\frac{S_{lm} - (-1)^m \bar{S}_{lm}}{2\eta\sqrt{l(l+1)}} \tag{57}
$$

with an effective torque and tension density resulting from the change in viscous forces given by

$$
\begin{aligned}
\frac{S_{lm}}{2} = \sum_{l_1,l_2,m_1,m_2} & \frac{(-1)^m \chi_{l_1 m_1}\delta R_{l_2 m_2}/R}{R^2/l_h^2 + (\nu+1)l_1(l_1+1) - 2} \\
& \sqrt{\frac{(2l_1+1)(2l_2+1)(2l+1)}{4\pi}} \begin{pmatrix} l_1 & l_2 & l \\ m_1 & m_2 & m \end{pmatrix} \\
& \left\{ \left[l_2(l_2+1) - 2\right]\sqrt{l_1(l_1+1)} \begin{pmatrix} l_1 & l_2 & l \\ -1 & 0 & 1 \end{pmatrix} \right. \\
& \left. - (\nu+1)l_1(l_1+1)\sqrt{l(l+1)} \begin{pmatrix} l_1 & l_2 & l \\ 0 & 0 & 0 \end{pmatrix} \right\}.
\end{aligned} \tag{58}
$$

Here

$$
\begin{pmatrix} l_1 & l_2 & l \\ m_1 & m_2 & m \end{pmatrix} \tag{59}
$$

are Wigner 3j symbols (closely related to the better known Clebsch-Gordan coefficients). They result from the product of two spherical harmonics function projected onto a third spherical harmonic, corresponding to the flow field that results from the couple of deformation and contractility fields. Details of the calculation will be published elsewhere. For small friction, i.e. large hydrodynamic length $l_h > R$, the deformation-dependent flow field resulting from $\delta A_{lm}, \delta B_{lm}$ is dominated by the deformation-triggered rotation given by $\delta B_{1,m}$. Identifying the rotation axis as the $z$ axis such that $\delta\mathbf{\Omega} = \mathbf{z}\delta\Omega$, the rigid body rotation of the deformed sphere is given by

$$\delta\Omega = -\sqrt{\frac{3}{4\pi}}\delta B_{1,0} = i\frac{3}{4\pi\gamma}\sum_{l,m}(-1)^m \chi_{l,m}\delta R_{l,-m}\frac{m[l(l+1)-2)]}{(\nu+1)l(l+1)-2}. \tag{60}$$

Writing the spherical harmonics coefficients in terms of an azimuthal angle and a magnitude as

$$\delta R_{lm} = |\delta R_{lm}|e^{-im\varphi_R}, \quad \delta\chi_{lm} = |\delta\chi_{lm}|e^{-im\varphi_\chi}, \tag{61}$$

we obtain

$$\delta\Omega = \frac{1}{\gamma}\sum_{l\geq 2,m}\frac{3}{4\pi}\frac{m\,[l(l+1)-2]}{(1+\nu)l(l+1)-2}|\chi_{l,m}||\delta R_{l,m}|\sin[m(\varphi_\chi - \varphi_R)] \tag{62}$$

This equation is equivalent to Eq. 45. We observe again that the rotation results from a misalignment between the heterogeneity of the surface geometry ($\delta R$) and the normal force pattern which we can express here directly in terms of the active contractility $\chi$. We observe that $\delta\Omega \to 0$ for $\nu \to \infty$, even if $\nu\eta =$ const., corresponding to vanishing shear viscosity. For vanishing bulk viscosity $\nu\eta \to 0$, $\delta\Omega$ does not vanish. Hence, the deformation-triggered rotation of an active fluid driven purely by active contractility results from viscous shear forces.

### 3 Dynamics of axis alignment

In the following, we study the dynamics of the cell division axis resulting from the compression-triggered rotation, i.e.

$$\partial_t\varphi = \delta\Omega, \tag{63}$$

where $\varphi$ is the azimuthal angle of the spindle axis of the AB cell and $\delta\Omega$ is the compression-triggered rotation given in Eq. 45. We consider a compression given by

$$\delta X_n \approx d_2(s)\cos(2(\theta - \varphi_2^X)), \tag{64}$$

where $\varphi_2^X$ defines a compression axis that is consistent throughout the embryo, i.e. $\partial_s\varphi_2^X = 0$. Also for the normal force pattern we consider a common axis throughout the embryo, which corresponds to the cell division axis, i.e. $\varphi_2^f(s) = \varphi$. In the following, we choose a coordinate system such that $\varphi_2^X = 0$. With this, the dynamics of the spindle axis can be written as

$$\Omega = \partial_t\varphi(t) = -W(t)\sin 2\Delta\varphi(t), \tag{65}$$

where $W$ is the magnitude of an effective force that drives alignment and can be expressed in terms of deformation and egg-shell normal force as

$$W(t) = -\frac{\pi}{\gamma\Theta_{zz}} \int_0^L \mathrm{d}s \rho(s) f_2(s,t) d_2(s). \tag{66}$$

Indeed, we find experimentally that $|\Omega|$ generally increases up to an angle of $\phi = \pm 45°$, i.e. $\sin 2\phi = \pm 1$, with $\phi$ being the angle of the cytokinetic ring (Fig. S6F). $f_2$ is the $k = 2$ component of the normal force for an axisymmetric egg-shell as defined in Eq. 42. It is given by the normal force balance equation Eq. 9 with the flow field obeying the tangential force balance equation 8. Hence $f_2$, is linear in the magnitude of the active tension driving the flow and depends on the viscosity of cortex and cytoplasm. Due to this linearity and rotational symmetry of the surface, $f_2$ only depends on the $k = 2$ component of the active tension $\chi$, which we interpret as the active tension in the cytokinetic ring $T(t)$ such that we may write

$$W(t) = \alpha T(t) \tag{67}$$

with $\alpha$ being a constant of proportionality that depends on the viscosities as well as the AP profile (i.e. $s$ dependence) of deformation and active tension. Eq. 62 yields an explicit expression for $\alpha$ for a deformed sphere.

We note that Eq. 65 is not specific to the active fluid model we have discussed in the previous sections. It is a general result for a rotation of an axis, here division axis, driven by misalignment with an external axis, here compression axis in two dimensions. It is solved by

$$\tan\varphi(t) = \tan\varphi(t_0) \exp\left[-\int_{t_0}^{t} \mathrm{d}t W(t)\right], \tag{68}$$

i.e. a exponential decay of $\tan\varphi$ with a time-dependent decay rate $W(t)$. The physical mechanism driving the alignment defines the time-evolution of this rate. When alignment is driven by actomyosin contractility and contractility scales linearly with Myosin concentration, $W(t)$ is proportional to the $k = 2$ component of Myosin concentration as measured by flourescence microscopy, which we call the relative Myosin concentration $M$ in the cytokinetic ring, see Suppl. Methods for details. Experimentally, we find that the time-evolution of $M$ can be captured by an exponential growth with a common rate $\lambda = 1/(13\mathrm{s})$, consistent with an instability of the cortex triggered by spindle-cortex interactions. Hence, $W(t) \sim T(t) \sim M(t)$ yields

$$\log\tan\varphi(t) = \log\tan\varphi(t_0) - \alpha\frac{M(t) - M(t_0)}{\lambda}. \tag{69}$$

Indeed, the experimental trajectories of $\varphi$ collapse onto such a curve (Fig. 4,S6).

### Supplementary methods

#### Experimental Methods

*Animal strains and culturing*

*C. elegans* strains were cultured using standard culture conditions[1] and maintained at 20C, apart from strains carrying *nmy-2(ne3409ts)*, which were maintained at 15C. *C. elegans* alleles and transgenes used in this study are: LGI, *nmy-2(ne3409ts)*[2], *nmy-2(cp8[nmy-2::gfp])*[3], LGII, *lin-5(he330[lin-5::glo-epdz::mcherry])*[4], LGV, *ruIs57[GFP::tubulin]*, LGIV, *cxTi10816(he259[ph::co-egfp::co-lov])*[4], LG unknown, *gesIs003[lifeact::mkate2]*[5]. Mouse embryos were from a CD-1(ICR) background in natural mating (without hormone indication).

*RNA interference*

RNAi treatment was performed by feeding as previously described[6]. Briefly, NGM agar plates containing 1 mM isopropyl-β-D-thiogalactoside and 50 µg/ml ampicillin were seeded with bacteria expressing dsRNA targeting the gene of interest or containing empty vector (L4440). The L4440 control and *lin-5(RNAi)* clones were obtained from the Ahringer RNAi library (Source BioScience)[7]. L4 larvae were grown on feeding RNAi plates at 20C for 25-27 hrs.

*Microscopy setups*

All *C. elegans* imaging was done using spinning disc microscopy on a Zeiss Axio Observer Z1 inverted microscope equipped with a Yokogawa CSU-X1 scan head, a C-Apochromat 63X/1.2 106 NA W objective, a Hamamatsu ORCA-flash 4.0 camera, 488 and 560 lasers, and operated by Micromanager software. Imaging was performed at room temperature (22-23C) apart from the experiments using the *nmy-2(ne3409ts)* and the accompanying controls. For these experiments the Cherry temp microscopy stage (Cherry Biotech) was used to switch between 15C and 25C. Mouse DIC imaging was done using a Zeiss Axio Observer Z1 inverted stand equipped with DIC optics, an NA0.55 WD 26 mm condenser, a 40x/NA1.1W objective and a Zeiss Axiocam 705 MRm monochrome CCD camera. Temperature (37.5C) and CO2 levels (5%) and humidity were maintained using a stage top incubator. An objective heater was used to prevent heat dissipation.

*Mounting and image acquisition*

Worm embryos were dissected from young adults and mounted in M9. For mild compressions worms were mounted on 2% agarose pads. For stronger compressions, embryos were mounted in buffer containing 10 or 15 µm polystyrene spacer beads (Polysciences). Subsequently a glass slide was lowered over it such that the embryos were confined between the cover glass and the glass slide. The samples were sealed using valap to prevent evaporation of the buffer during imaging. In order to obtain uncompressed embryos, embryos were mounted in M9 on a coverslip coated with poly-L-lysin. Playing clay was fixed at the corners of the coverslip in order to achieve excess spacing.

For dual-color imaging of Lifeact-mKate2 and GFP-tubulin (control, non-RNAi and embryos carrying *nmy-2(ne3409ts)* without *opto-lin-5*) embryos were imaged using a 488 and 560 laser with 9-11 second time intervals. At each timepoint a z-stack was made of 26 slices (compressed embryos) or 31 slices (uncompressed embryos) with 1 um spacing. For single color imaging of GFP-tubulin, embryos were imaged with 8 second time intervals (*lin-5(RNAi)*, *opto-lin-5* and accompanying controls) or 10 second time intervals (*opto-lin-5; nmy-*

*2(ne3409ts)* and accompanying controls). At each timepoint a z-stack was made of 26 slices with 1 um spacing. For the experiments using the *nmy-2(ne3409ts)* allele and the accompanying controls, embryos were kept at 15C until 3-4 min after completion of the first cytokinesis. Thereafter, the temperature was switched to 25C, followed by the start of image acquisition.

For imaging of the actomyosin cortex with high time resolution (dt=2 seconds), embryos producing endogenously labeled NMY-2::GFP (*nmy-2(cp8[nmy-2::gfp])*) were used. Every time point (dt=2 sec) 2 focal planes with a z-spacing of 1 um were recorded, as well as a midplane slice 10 um above the cortical plane. For analysis a maximum intensity projection was made of the 2 cortical slices. Before the time lapse acquisition, a single z-stack was recorded of 31 slices with 1 um spacing to extract the embryo outline in the DV-LR plane. For mouse zygote imaging, mouse cumulus complexes, including zygotes, were isolated from the oviduct of donor animals after a plug check at 8:00 am. With the help of the enzyme hyaluronidase, zygotes were washed out of the cumulus complex using M2 medium and selected for the presence of pronuclei. Just before the experiment, zygotes were placed in glass capillaries (Hirschmann Ringcaps 50µl, handpulled, inner diameter: 60-65µm) using a mouth pipette. These were then transferred to a 200µl droplet of KSOM media in dishes for incubation and microscopy (MatTek Corporation P35G-0.170-14-C), overlaid with Paraffin (Sigma-Aldrich, 1.07160.1000), and incubated and imaged at 37 degrees with 5% CO2. To generate the time-lapse recordings, every 60 seconds a 60-um z-stack with 2 um z-spacing was acquired. Subsequently, for every time point the midplane z-slice was extracted.

**Image pre-processing and analysis**

For visualization and analysis of the DV-LR plane, the image stacks were first rotated to align the AP axis horizontally. Images from the stacks were then cropped in the AP-DV plane to a 150-pixel wide region (15.87 um) that spans the central part of the AB cell. Subsequently, a projection along the z axis, i.e. the AP axis, was made in every timepoint and pixel values were interpolated on a grid with 0.1058 um spacing (corresponding to the pixel size in the x-y plane, i.e. the AP-DV plane) using cubic interpolation.

Spindle length (pole-to-pole distance) and angle with the DV axis were extracted by manually tracking the spindle poles, using the GFP-tubulin channel, in the DV-LR plane. To compute averages and standard deviations of the spindle orientation across embryos, we first computed an average phase factor corresponding to an average nematic order parameter as

$$Q_{av} = \langle e^{2i\phi} \rangle$$

with $\phi$ being the spindle angle. The average angle $\phi_{av}$ and its (circular) standard deviation $\phi_{std}$ were then computed as

$$\phi_{av} = (arg\ Q_{av})/2, \quad \phi_{std} = \sqrt{-\,log|Q_{av}|\,/2}$$

The AB cell outline in the DV-LR plane was extracted from the Lifeact-mKate channel or the NMY-2::GFP channel by applying segmentation using an intensity threshold. Because Lifeact-mKate levels varied among embryos the segmentation was manually supervised by varying the intensity threshold. Aspect ratios in the DV-LR plane were derived from these outlines by dividing the LR distance by the DV distance.

To extract the position of the cytokinetic ring from the Lifeact-mKate channel, a maximum intensity projection was first made of the two z-slices (dz=1 um), closest to the imaging objective, that were cropped to a 30-pixel wide region centered at the center of the AB cell in the AP-DV plane. For every row of pixel values, the mean pixel values were extracted, yielding a vertical line of mean pixel values along the DV axis in each time point. Subsequently, these were concatenated to make kymographs from which the intensity of the cytokinetic ring was extracted. In addition, together with the AB cell outline in the DV-LR plane, this kymograph was used to extract the orientation of the cytokinetic ring in the DV-LR plane (see below).

The end point of the AB rotation was defined as the onset of the cell division skew on the AP-DV plane (Fig S1C). For embryos producing a cortical marker (lifeact-mKate2 or NMY-2::GFP) the onset of rotation was defined as the first time point in which rotation was observed. For embryos without a cortical marker, the time point 160 seconds prior to the end point was taken as the first time point of analysis.

In order to determine the orientation of the ring, we assigned an azimuthal angle to each pixel of the outline of the cortex in the DV-LR plane. To do so, we used the center of mass $\mathrm{r}_0 = (\mathrm{x}_0, \mathrm{y}_0) = \langle(\mathrm{x}_\mathrm{j}, \mathrm{y}_\mathrm{j})\rangle$, determined from the positions $\mathrm{r}_\mathrm{j} = (\mathrm{x}_\mathrm{j}, \mathrm{y}_\mathrm{j})$ of the pixels belonging to the cortex. Here x- and y-axis correspond to the axes in the LR-DV plane that are perpendicular and parallel to the imaging plane respectively. Due to compression perpendicular to the imaging plane the x (y)-axis corresponds to the short (long) axis in the LR-DV plane. The azimuthal angle $\theta_\mathrm{j}$ for each pixel j in the LR-DV plane was then determined using

$$(x_j - x_0) + \mathrm{i}(y_j - y_0) = r_j exp(i\theta_\mathrm{j}) \qquad (S1)$$

Where i is the imaginary unit and $r_j > 0$. With this we fitted the function

$$\mathrm{f}(\theta) = f_0 + f_1 cos2\theta + f_2 sin2\theta \qquad (\mathrm{S2})$$

to each time-point of the kymograph consisting of measured intensities $I_j$ of LifeAct-mKate in the cortical plane. For each time point, we define the magnitude or ring intensity M and angle $\phi$ of the emerging cytokinetic ring as

$$f_1 + \mathrm{i}f_2 = \mathrm{M}\exp(2i\phi) \qquad (S3)$$

where $\mathrm{M} > 0$, such that $\phi = 0$ corresponds to a ring aligned with the x-axis, i.e. the short axis in the LR-DV plane (See Fig. S6A for an example). In Fig. 2D, the ring intensity M was normalized using the maximum value of M within the time interval of rotation for each embryo.

For quantifying movements of the actomyosin cortex in the AB cell with high time resolution (dt=2s), Particle Image Velocimetry (PIV) was performed using an open source Matlab package (PIVlab)[8]. A 3-step PIV with a final box size of 36x36 pixels, on a grid with 18 pixel spacing, was performed using a manually generated mask that segments the cortex of the AB cell in the AP-DV plane. The obtained flow fields were rotated to align the anteroposterior axis horizontally (anterior left, posterior right). Subsequently, velocity vectors at the anterior and posterior-most 15% of the AB cell mask were excluded. Of the remaining velocity vectors, the mean of the components perpendicular to the AP axis (DV velocity) was computed every time point. To visualize the movements of the AB cell together with the P1 cell, PIV was performed, with the same settings, on a movie rotated such that the AP axis is horizontally oriented. The analysis was done using a manually generated mask that segments the cortical of both the AB cell and the P1 cell. Cortical flow vectors were averaged over the time period of the rotation, to obtain the mean flow field.

To extract the position of the cytokinetic ring, and the ring intensity over time, from the cortical NMY-2::GFP signal, movies were first rotated to align the AP axis horizontally. Subsequently, a 90-pixel wide region of interest, covering the center part of the AB cell in the AP-DV plane, was used to make the kymograph. For every row of pixel values, the mean pixel values were extracted, yielding a vertical line of mean pixel values along the DV axis in each time point. Subsequently, these were horizontally concatenated to make kymographs from which the position and intensity of the cytokinetic ring were extracted.

For the last time point of the rotation we determine the orientation $\phi_N$ of the ring from the kymograph as described above. For earlier time points, however we make use of the average DV velocity $v_{DV}$ as measured by PIV. From this, we compute an angular velocity $\Omega = v_{DV}/\mathrm{R}$ where R is the average radius of the outline of the cortex determined from the cross-section in the DV-LR plane. Then, the orientation $\phi_j$ at time point j is computed as

$$\phi_j = \phi_N - \sum_{k=j}^{N} \Omega_k \times 2\mathrm{s}, \qquad (S4)$$

Where $\Omega_k$ is the angular velocity between time points $\mathrm{k}$ and $\mathrm{k}+1$. Thereby, we circumvent that the ring is often found outside of the cortical plane for early time points which can make the orientation determined from intensity measurements in the cortical plane unreliable (See Fig. S6B). This method is based on the observation that movements of cortex as well as cytoplasm resemble the rotation of a rigid body, such that the rotational velocity of the cortical plane equals the rotational velocity of the cytokinetic ring independent of the position of the cytokinetic ring. The ring intensity $M_{raw}$was determined using linear regression, fitting the function

$$\mathrm{f}(\theta) = f_0 + M_{raw} cos\left(2\theta - \phi_j\right) \qquad (S5)$$

to the intensity values of time point j from the cortical kymograph. Also in the absence of a cytokinetic ring, NMY-2:GFP intensities are never uniform in the cortex, which often yields negative vales for $M_{raw}$ for early time points (Fig. S5A). These inhomogeneities and the resulting $M_{raw}$ at early time points differ in position and magnitude between embryos. However, we find that the trajectories of $M_{raw}$ collapse onto an exponential curve $exp[\lambda t]$ with a common growth rate $\lambda = 1/(13s)$ for early time points, when allowing for a constant embryo-specific offset (Fig. S5E). The offset $M_0$ is determined by fitting the function

$$\mathrm{f}(t) = M_0 + \mathrm{A}exp[t/13s] \qquad (S6)$$

to $M_{raw}$ for each embryo using linear regression. With this we define the ring intensity

$$\mathrm{M} = M_{raw} - M_0 \qquad (S7)$$

which we use in Fig. 3B. Since the ring intensity follows an exponential growth, our coarse-grained model predicts that the ring angle $\phi$ can be written as

$$\phi(t) = \arctan\left[tan\phi(0)exp\left[-\alpha/\lambda\left(M - M(0)\right)\right]\right], \qquad (S8)$$

if the effective ring tension driving alignment is proportional to $\mathrm{M}$ (See. Eq. 69 in supplementary notes). Here $\alpha$ is a constant that depends on the geometry and the effective viscosity of the embryo. We consider equal values of $\alpha$ for all embryos. With this, the model predicts that $\phi - \phi_0$ is a common function (Eq. S8) of $M - M(\phi(t) = \phi_0) = M_{raw} - M_{raw}(\phi(t) = \phi_0)$. Here $\phi_0$ is an arbitrary reference angle. In Fig. S5G-H, we use $\phi_0 = 30°$ and define $M_{raw,0} = M_{raw}(\phi(t) = \phi_0)$. We find that the trajectories do indeed collapse onto a common function given by Eq. S8. $\alpha$ is determined by fitting a linear curve to $log\, tan\, \phi(t)$.

To measure cytoplasmic flow in the AB cell in the DV-LR plane, Particle Image Velocimetry (PIV) was performed on the lifeact-mKate2 signal in the time window spanning the AB rotation. First, a mask segmenting the cytoplasm and excluding the cortical signal lifeact-mKate2 signal was generated manually. Subsequently, 3-step PIV[8] was performed with final box size of 22x22 pixels (step-size of 12 pixels), on a grid with 11 pixel spacing. Cytoplasmic flow vectors were averaged over time to generate a mean cytoplasmic flow field. For visualization the DV component (y-component) of the mean flow vectors was color-coded.
